# mini-MEndR: A 96-well mixed species culture assay to co-evaluate human and mouse Pax7^+^ cell-mediated skeletal muscle repair

**DOI:** 10.1101/2023.01.20.524941

**Authors:** Nitya Gulati, Sadegh Davoudi, Bin Xu, Saifedine T. Rjaibi, Erik Jacques, Justin Pham, Amir Fard, Alison P. McGuigan, Penney M. Gilbert

## Abstract

Functional evaluation of molecules that are predicted to promote stem cell mediated endogenous repair often requires *in vivo* transplant studies that are low throughput and hinder the rate of discovery. Here, we offer a strategy to rapidly test and prioritize molecules for functional validation studies. We miniaturized, simplified and expanded the functionality of a previously developed muscle endogenous repair (MEndR) *in vitro* assay that was shown to capture significant events of the first week of the *in vivo* muscle endogenous repair process. The new “mini-MEndR assay” consists of miniaturized cellulose scaffolds designed to fit in 96-well plates. The scaffold pores are infiltrated with myoblasts encapsulated in a fibrin-based hydrogel to form thin, engineered skeletal muscle tissues. By evaluating multiple commercially available human primary myoblast lines in 2D and 3D culture, we establish quality assurance metrics for cell line selection that standardize myotube template quality. Pre-adsorbing thrombin to the cellulose scaffolds facilitates *in situ* tissue polymerization, a critical modification that enables users proficient in myoblast culture to rapidly acquire myotube template fabrication expertise. Following the generation of the 3D myotube template, muscle stem cells (MuSCs), enriched from digested mouse skeletal muscle tissue using an improved magnetic-activated cell sorting protocol, are engrafted onto the engineered human muscle template. A regenerative milieu is then introduced by injuring the muscle tissue with a myotoxin. Addition of a known modulator of MuSC mediated repair recapitulates the *in vivo* outcomes (enhanced muscle production and Pax7^+^ cell expansion), but only in the presence of both the stem cells and the regenerative milieu. By fluorescently labeling the mouse MuSCs, we demonstrate the feasibility of co-evaluating human and mouse Pax7^+^ cell responses to drug treatment, thereby expanding the utility of the assay. Importantly, phenotypic data is collected with a high-content imaging system and is analyzed using CellProfiler-based image analysis pipelines. The miniaturized predictive assay offers a simple, scaled platform with which to co-investigate human and mouse skeletal muscle endogenous repair molecular modulators, and thus is a promising strategy to accelerate the muscle endogenous repair discovery pipeline.

## Introduction

The regenerative capacity of healthy skeletal muscle is driven by the activity of tissue resident muscle stem cells (MuSCs). With age^1^, or in disease, e.g. Duchenne muscular dystrophy (DMD)^2,3^, MuSCs remain present, but have diminished repair capabilities. Therapeutic strategies to correct diminished skeletal muscle endogenous repair have a growing need. One exciting emerging strategy to improve skeletal muscle endogenous repair is the use of drugs to boost MuSC function (i.e. endogenous repair), for which pre-clinical studies in mice demonstrate the promise of this therapeutic approach^3–6^.

A current gold standard assay to determine whether a molecular target restores MuSC function, and thus to make a go/no-go decision in the discovery process, is the intramuscular transplantation assay^7–9^. This assay involves transplanting treated cells into the pre-irradiated and pre-injured hindlimb of immune-compromised rodents and then assessing the capacity of the transplanted cell population to make muscle and repopulate the stem cell compartment. This gold-standard assay is expensive, low throughput and plagued by high variation, which in turn limits the number of candidates that can be tested. To overcome this pain point in the discovery process, we^10^ and others^11,12^, have invented in vitro functional assays to predict skeletal muscle endogenous repair outcomes “in a dish” as a lower cost complement to the intramuscular transplantation assay. Most of the assays evaluate engineered skeletal muscle microtissue repair after an induced injury. Since the primary myoblast cultures used to produce the microtissues contain Pax7^+^ cells, tissue repair is attributed to the activity of this sub-population. By contrast, the muscle endogenous repair (MEndR) assay tracks the activity of bonefide MuSCs engrafted into thin engineered muscle tissues that are suited to long-term timelapse microscopy^10^.

In the MEndR culture assay, we first engineer thin sheets of mature skeletal muscle tissue by infiltrating a thin cellulose (paper) scaffold with primary patient-derived muscle progenitor cells encapsulated in an extracellular matrix hydrogel. MuSCs freshly isolated from mouse hindlimb skeletal muscle tissue are labelled and then incorporated into the 3D muscle sheets. Shortly thereafter, the muscle tissue sheet is injured by exposure to a myotoxin to induce a regenerative microenvironment, and MuSC-mediated muscle tissue template repair is assessed phenotypically by assessing the contribution of labelled MuSCs to new muscle tissue formation, and to niche repopulation, using fluorescent image analysis. To validate the predictability of our platform, we tested a number of drugs that were previously reported modulators of MuSC function and compared results using MEndR with those we obtained in parallel using the *in vivo* transplantation assay. MEndR accurately predicted *in vivo* transplantation assay outcomes. Side-by-side comparisons accurately identified a range of repair phenotypes observed in the *in vivo* transplantation assay results. Furthermore, since MEndR tissues are thin and flat, the user has greater control over the conditions during drug testing than with the previously described microtissue based assays^11,12^ (e.g., MuSC content, state of the template, uniform exposure to media components). With this prior study^10^, we demonstrated for the first time, the ability to compress the translational pipeline by offering a complement to the *in vivo* validation assay that is, in theory, amenable to testing all 2D drug screen ‘hits’ without the need for biased pre-selection.

The MEndR functional assay is poised to serve as a secondary screen service to prioritize ‘hits’ arising from high-throughput screens designed to discover MuSC potency modulators. However, MEndR assay scale-up from a 24-well footprint to a 96-well format is critical to realizing this goal. Candidate assessment performed in a 96-well format has the potential to offer an order of magnitude higher throughput than the intramuscular transplantation assay. Towards this goal, we herein describe the scaling of the MEndR assay into a 96-well footprint and we validate that the miniaturized platform captures all stem cell functional metrics shown in the original MEndR platform. We also extend the utility of the assay by demonstrating the capacity to co-evaluate responses of human (within myotube template) and mouse (engrafted MuSCs) Pax7^+^ populations to molecule treatments.

## Materials and Methods

### Human myoblast maintenance and 2D differentiation

Human primary myoblasts (pMB) were purchased from Cook MyoSite^®^ (Table 1). For routine culture, myoblasts were plated on collagen-coated tissue culture plates (Table 2) at a seeding density of 5000 - 7500 cells/cm^2^. Myoblasts were maintained in human pMB growth medium (GM; Table 2) and passaged using trypsin upon reaching ∼70% confluency. Culture media was replaced every other day. Myoblasts were used at passage 6-7 for 2D experiments, and at passage 8 in all 3D tissue experiments. For 2D differentiation studies, myoblasts were plated at a seeding density of ∼37,000 cells/cm^2^ on Geltrex^TM^ coated plates in human pMB GM. The cultures were transitioned to 2D differentiation media (DM; Table 2) one-day post seeding to induce fusion. Half of the DM was removed and replaced every other day. Cultures were fixed and analyzed on day 9 of differentiation.

**Table 1.** Human skeletal muscle myoblast donor information.

| Age | Sex | Ethnicity | Vendor | Catalog # | Tissue of Origin | Additional Information |
| --- | --- | --- | --- | --- | --- | --- |
| 18 | M | Caucasian | Cook MyoSite | SK-1111-P01236-18M | Rectus Abdominus | Active |
| 18 | F | Caucasian | Cook MyoSite | SK-1111-P01431-18F | Vastus Lateralis | Healthy, No known medical conditions |
| 19 | F | Caucasian | Cook MyoSite | SK-1111-P01358-19F | Vastus Lateralis | N/A |

**Table 2.** Culture medium and solutions.

| Media | Composition |
| --- | --- |
| Wash Medium | 10 % FBS, 90 % DMEM |
| Human pMB Growth Medium | Ham's F-10 nutrient mix, 20 % Foetal bovine serum FBS, 5 ng/mL rh-FGF2, 1 % Penicillin-Streptomycin |
| 2D Differentiation Media | DMEM, 2 % Horse serum, 10 g/mL insulin (Sigma, #I6634), 1 % Penicillin-Streptomycin |
| Rat Tail Collagen Solution | 0.1 M Acetic Acid, Collagen I Rat Tail Stock solution (Gibco #A1048301), 0.1 M Acetic Acid, Collagen Stock Solution |
| MuSC Growth Medium | DMEM/F12, 1 % Pen-strep (P/S), 20 % Foetal bovine serum (FBS), 10 % Horse serum (HS), 1 % 2 mM Q (Glutamax), 1 % Insulin-Transferrin-Selenium (ITS), 1 % Non-Essential Amino acids, 1 % 1 mM Sodium Pyruvate, 50 uM (1/1,000) β-mercaptoethanol, 5 ng/ml (1/5,000) Gibco FGF2 |
| MEndR Growth Medium | Ham's F-10 nutrient mix, 20 % FBS, 1.5 mg/mL 6-Aminocaproic Acid (ACA, Sigma, #A2504), 1 % Penicillin-Streptomycin |
| MEndR Differentiation Medium | DMEM, 2 % Horse serum, 2 mg/mL ACA, 10 g/mL insulin (Sigma, #I6634), 1 % Penicillin-Streptomycin |
| Fibrinogen Solution | 10 mg/mL Fibrinogen (Sigma, #F8630) in NaCl solution (0.9 % wt/vol in ddH<sub>2</sub>O) |
| Extracellular Matrix (ECM) Master Mix | 40 % DMEM, 40 % Fibrinogen, 20 % Geltrex™ |
| Red Blood Cell Lysis Buffer | H<sub>2</sub>O, 0.155 M NH<sub>4</sub>Cl, 0.01 M KHCO<sub>3</sub>, 0.1 mM EDTA |
| FACS Buffer | PBS, 2.5 % goat serum, 2 mM EDTA |
| Blocking Solution | PBS, 10 % goat serum, 0.3 % Triton X-100 (BioShop, #TRX777) |

### MEndR and mini-MEndR myotube template fabrication and injury

24-well plate format myotube templates were generated as previously described by our group^10^. Briefly, rectangle patterns with 0.5 cm x 1 cm dimensions were printed onto the cellulose scaffold product (MiniMinit® Products Ltd) with a LaserJet printer and then cut into individual rectangular scaffolds with scissors. These scaffolds were then autoclaved in a dry cycle to maintain sterility. To generate the myotube templates, sterile cellulose scaffolds were carefully placed on a custom polydimethylsiloxane (PDMS) block using a pair of sterile tweezers. The PDMS block was placed in an ethanol-sterilized plastic container with moist paper on the bottom and a tight-fitting lid to seal the top, to create a humidified chamber. Next, a fibrinogen solution (Table 2) was freshly prepared and filter-sterilized with a 0.22 µm filter. pMBs were resuspended at a concentration of (3.33E+07 cells/mL) in an extracellular matrix (ECM) master mix (Table 2). 100 U/mL of thrombin (Sigma, #T6884) was added to the pMB/ECM mixture to achieve a final concentration of 0.225 U/mg of fibrinogen. The mixture was maintained on ice throughout. Next, 12 µL of pMB/ECM mixture (i.e., 4.0E+05 pMBs) was evenly distributed onto the cellulose scaffolds using a custom spreading tool^13^. The humidified chamber was then sealed with the lid and incubated at 37 °C for 5 minutes to encourage conversion of fibrinogen to fibrin clots. Next, MEndR GM (Table 2) was gently pipetted onto each of the scaffolds to facilitate easy detachment from the PDMS block. The scaffolds were then gently lifted and placed into individual wells of a 24-well plate containing 1 mL of MEndR GM. Two days later, the GM was removed and replaced with MEndR DM to facilitate myotube differentiation. Half of the MEndR DM was removed and replaced every other day.

For 96-well plate format myotube templates (mini-myotube templates), cellulose paper (MiniMinit® Products Ltd) was cut into circular discs using a 5 mm biopsy punch (Integra, #MLT3335) and subsequently sterilized in an autoclave. 96-well plates (PhenoPlate 96-well microplates, PerkinElmer) were prepared for tissue seeding and culture by treatment with 100 μl of 5 % Pluronic F-127 (Sigma, Cat#P2443) solution in dH_2_O (Table 2) at 4 °C overnight, or minimum one hour, which served to create a non-adhesive surface to prevent cell attachment on the plates and facilitate tissue self-organization. The pluronic acid was then aspirated, and the plate was left to dry in the biosafety cabinet with the lid removed until the thin film of liquid could no longer be visualized (∼20 min). The autoclaved disks were then placed in the center of the corresponding wells of the 96-well plate using a sterile anti-static tweezer. Next, a 0.8 U/mL thrombin solution was prepared by diluting the 100 U/mL thrombin stock in PBS, and 4 µL of the 0.8 U/mL thrombin solution was dispensed onto the center of each paper disc. The discs were then left to dry in the biosafety cabinet. Meanwhile, a 10 mg/mL fibrinogen solution (Table 2) was freshly prepared by dissolving lyophilized fibrinogen powder in 0.9 % wt./vol of NaCl solution, and subsequently sterilized by passing through a 0.22 µm filter. pMBs were trypsinized, counted and suspended in ECM master mix (Table 2) at a concentration of 2.5E+07 cells/mL, which was kept on ice. Next, 4 µL of pMB/ECM mixture (i.e., 1.0E+05 pMBs) was pipetted into the center of paper discs with pre-adsorbed thrombin, with the mixture entering the cellulose scaffold pores by capillary action. Cellulose scaffolds impregnated with pMB/ECM were incubated for 5 minutes at 37 °C. 200 µL of MEndR GM (Table 2) was then gently pipetted into each well, ensuring that the tissues did not flip over. The 96-well plate was then returned to the cell culture incubator (37°C, 5 % CO_2_). The next day, tissues that failed to detach from the bottom of the well (by visual inspection), were released by gently triturating the culture media in the well with a p200 pipette. Two days later, the MEndR GM was aspirated and replaced with MEndR DM (Table 2). Half of the MEndR DM was removed and replaced with fresh medium every other day.

### Immunohistochemistry

2D myotube cultures, and 3D muscle tissues, were harvested and fixed at the indicated culture endpoints by removing the media, washing gently three times with phosphate buffered saline (PBS), and then incubating in 200 µL of 4% paraformaldehyde for 15 min at room temperature. The samples were washed three times with PBS every 5 min followed by incubation in blocking solution (Table 3) overnight at 4 °C, or for 1 hr at room temperature. Following blocking, which also contains permeabilization agent, samples were then incubated with 40 µL of primary antibodies (Table 3) diluted in PBS and 0.1 % goat serum, overnight at 4 °C. Next, the samples were washed three times with cold PBS every 10 min and then incubated for 45-60 min at room temperature with 40 µL of a secondary antibody mix (Table 3), containing DRAQ5 nuclear counter stain (Table 3), and diluted in PBS with 0.1% goat serum. The secondary antibody mix contained phalloidin conjugates (Table 3) for the experiments where phalloidin coverage was assessed. The samples were then washed three times with cold PBS. Tissue samples were stored in PBS at 4 °C until image acquisition.

**Table 3.** Antibodies and other reagents.

| Antibody | Host species | Dilution | Manufacturer |
| --- | --- | --- | --- |
| Alexafluor® 647 Anti-Human CD56 | Mouse | 1:20 (FC) | BD, #557711 |
| eFluor® 660 Anti-Mouse CD34 | Rat | 1:66 (FC) | eBioscience, #50034182 |
| PE Anti-Mouse α7-Integrin | Rat | 1:500 (FC) | AbLab, #530010-05 |
| Biotin Anti-Mouse CD31 | Rat | 1:200 (FC) | BD, #553371 |
| Biotin Anti-Mouse CD45 | Rat | 1:500 (FC) | BD, #553078 |
| Biotin Anti-Mouse CD11b | Rat | 1:200 (FC) | BD, #553309) |
| Biotin Anti-Mouse Ly-6A/E | Rat | 1:200 (FC) | BD, #553334 |
| Propidium Iodide | - | 1:1000 | Sigma, # P4864 |
| Anti-Sarcomeric α-actinin | Mouse | 1:500 | Sigma, #A7811 |
| Anti-GFP | Rabbit | 1:500 (IF) | ThermoFisher, #A11122 |
| Anti-Pax7 | Mouse | 1.5:1 (IF) | Iscove |
| Alexafluor® 488 Anti-Rabbit | Goat | 1:500 (IF) | ThermoFisher, #A11008 |
| Alexafluor® 546 Anti-Mouse | Goat | 1:500 (IF) | ThermoFisher, #A11003 |
| Alexafluor® 647 Anti-Rat | Goat | 1:500 (IF) | ThermoFisher, #A21247 |
| Phalloidin 568 | - | 1:500 (IF) | Life Technologies, #A12380 |
| Hoechst | - | 1:1000 (IF) | ThermoFisher, #H3570 |
| DRAQ5 | - | 1:1000 (IF) | Cell Signaling Technology, #4084L |
| DAPI | - | 1:1000 (IF) | Roche, #10236276001 |
| Satellite Cell Isolation Kit | Mouse | 1:5 (MACS) | Miltenyi Biotec, #130-104-268 |
| Anti-Integrin-α7 Beads | Mouse | 1:5 (MACS) | Miltenyi Biotec, #130-104-261 |
| Click-iT™ Plus EdU Cell Proliferation Kit | | (EdU, 1 μM) | ThermoFisher, #C10638 |

### Image acquisition

Confocal z-stack images of immunostained samples were acquired using an Olympus FV-1000 confocal microscope and Olympus FluoView V4.2b imaging software. Images were acquired using x4 (0.13 NA), x10 (0.3 NA) or x20 (0.45 NA) air objectives (Olympus) and the step size for each z-stack image acquired was set at the optimal value for each objective by the imaging software. Laser power, PMT HV and background offset were adjusted using the grey scale scan. Laser power was set lower than 75 % during image acquisition to prevent photobleaching and reduce cytotoxicity for all images. Next, laser power and HV were adjusted to avoid overexposure and saturation as guided by the red pixels. Background offset was adjusted to preclude blue pixels in each of the stacks. HV was set lower than 750 to reduce noise and Gain was set at 1 for all images. For the niche repopulation experiments, tissues were imaged using an Opera Phenix High Content Screening system outfitted with a x20 water immersion objective. Confocal stacks were converted to the 2D space using a maximum Z-projection prior to image analysis. Nuclear fusion index, Pax7^+^ cell enumerations, as well as sarcomeric α-actinin (SAA), green or yellow fluorescent protein (GFP/YFP) or phalloidin tissue coverage were quantified exactly as we previously described^10^.

### Bead retention assay and analysis

Cellulose-based paper scaffolds (MiniMinit® Products Ltd) were cut using a 5 mm diameter biopsy puncher and placed in a 96-well plate. Scaffolds were pre-adsorbed with thrombin at a concentration of 0.8 U/ml (0.2 U/mg fibrinogen) and allowed to dry for one hour in the biological safety cabinet with the 96-well plate lid removed. Next, a Master Mix was prepared by diluting 15 μm diameter fluorescent beads (Bangslab, #FCDG011) 10 times in ECM master mix (Table 2). Then, 4 μL of the bead MasterMix was infiltrated into each of the dried scaffolds and the scaffolds were subsequently incubated at 37 °C for 5-10 min to allow the Master Mix and thrombin to gel.

The scaffolds were imaged using confocal microscopy (Olympus FV-1000, x4 air objective, laser power 20-40 %, HV < 700) to determine the initial number of beads. Scaffolds were maintained in PBS while not being imaged to prevent them from drying. After the initial images were captured, scaffolds were washed with PBS three times to mimic the turbulence experienced by scaffolds during 3D culture (e.g., media changes). The scaffolds were imaged once more, using the same settings, to determine the number of beads retained.

To quantify the number of beads in an image, confocal stacks were first converted to the 2D space using a maximum Z-projection. Thresholding was performed using FIJI’s built-in Yen algorithm to binarize the image into fluorescent beads and background. Overlapping adjacent beads were segmented using the watershed tool. Finally, FIJI’s *Analyze Particles* function was used to count the number of beads recorded in the image.

### Mini-myotube template endogenous repair

To study myotube template endogenous repair potential, we optimized an injury protocol that achieved an ∼50 % reduction in SAA coverage of the myotube template post injury. Specifically, the culture media was removed from the myotube template tissues and replaced with MEndR DM containing 0.5 µM cardiotoxin (CTX) (Latoxan #L8102 – diluted in water) on day 7 of differentiation. Tissues were incubated on an orbital shaker in a 37 °C incubator for 4 hours. The medium was gently pipetted at the 2-hour mark to encourage the movement of the medium in the well and the detachment of any broken fibers. After the 4-hour incubation, cardiotoxin was removed by gently washing the tissues twice with MEndR DM. Tissues were then cultured in MEndR DM supplemented with 5 ng/mL rh-FGF2 for 24 hr, to assess template injury.

For myotube template self-repair studies, tissues were first cultured for a period of 5 days post injury, during which time the tissues were treated with endogenous repair stimulating drugs as indicated in Table 4. The MEndR DM supplemented with FGF2 and drugs was completely replaced every other day. For the remaining duration of the assay, tissues were cultured in MEndR DM and half of the media was replaced every other day.

**Table 4.** Small molecule information.

| Drug name | Target | Working concentration | Manufacturer |
| --- | --- | --- | --- |
| SB203580 | p38α/$\beta$ MAPK | 10 μM | New England biolabs, #5633 |
| Erlotinib Hydrochloride | Epidermal Growth Factor Receptor (EGF) | 5 μM | Sigma Aldrich, #sc-202154 |
| DMSO | - | - | Sigma Aldrich, #D2650 |

### Mouse strains and ethics

All animal use protocols were reviewed and approved by the local Animal Care Committee (ACC) within the Division of Comparative Medicine (DCM) at the University of Toronto. All methods in this study were conducted as described in the approved animal use protocols (#20012838) and more broadly in accordance with the guidelines and regulations of the DCM ACC and the Canadian Council on Animal Care. 129-Tg(CAG-EYFP)/7AC5Nagy/J mice are maintained by crossing homozygous males and females. C57BL/6-Tg (CAG-EGFP)1Osb/J (Actin-eGFP) mice were purchased from the Jackson Laboratory (USA) and utilized in experiments after a 1-week acclimatization period. Tg(Pax7-nGFP) mice^14^ were a kind gift from Dr. Shahragim Tajbakhsh, transferred from the laboratory of Dr. Michael Rudnicki and maintained at the University of Toronto by backcrossing to C57Bl/10ScSn (Jackson Laboratory) and genotyping for the GFP gene. For all experiments, 8-12 week old female mice were used.

### Murine muscle stem cell (MuSC) isolation

MuSCs were prospectively isolated from the collective mouse hindlimb muscles using Fluorescence Activated Cell Sorting (FACS) as previously described^15^. Briefly, mice were humanely euthanized, and their hindlimb muscles were removed and placed in a GentleMACS dissociation tube (Miltenyi Biotec, #130-096-334) containing Type 1A Collagenase (628 U/mL, Sigma, #C9891) in DMEM. The tissue was then physically dissociated by running the “Skeletal Muscle” program in the GentleMACS Dissociator (Miltenyi Biotec, #130-093-235). Next, the tube was placed on a rotating mixer at 37 °C for 1 hr. Dispase II (0.04 U/mL, Life Technologies, #17105041) was then added to the tube and the muscle was further dissociated by passing through a 10 mL pipette 10 times. Afterwards, the tube was returned to the rotating mixer for 1 hr. Next, the digested muscle was passed through a 21 G needle 10 times, after which 10 mL of FACS buffer was added to the tube. The digested muscle was passed through a 70 µm cell strainer, followed by a 40 µm cell strainer. The resulting cell slurry was centrifuged at 400 g for 15 min, after which the pellet was resuspended in 1 mL of FACS buffer. Red blood cells were removed by resuspending the cell pellet in red blood cell lysis buffer and incubating at room temperature for 7 min. The cells were then washed with FACS buffer and centrifuged at 400 g for 15 min.

Next, the cells were resuspended in 1 mL FACS buffer and incubated with the following antibodies for 1 hr on a rotating mixer at 4 °C: CD34-eFluor 660 (1:66), α7-integrin-PE (1:500), biotinylated CD31 (1:200), CD11b (1:200), Ly6A/E (1:200), and CD45 (1:500). The cells were then washed with FACS buffer, centrifuged at 400 g for 15 min, and resuspended in 1 mL FACS buffer. Streptavidin-PE-Cy7 (1:200) was added to the cell suspension, which was then placed on a rotating mixer at 4 °C for 30 min. The cells were then washed with FACS buffer a final time, centrifuged for 15 min at 400 g, and resuspended in 1 mL FACS buffer. Finally, propidium iodide (1:1000) was added to the cells, and MuSCs were enriched by FACS using the following gating strategy: CD34^+^/ITGA7^+^/CD45^-^/CD31^-^/CD11b^-^/Ly6A^-^/PI^-^.

The other method to isolate murine MuSCs from the collective mouse hindlimb muscles was to use Magnetic-Activated Cell Sorting (MACS) based on the vendor protocols (Miltenyi) with adaptations that we found to improve MuSC enrichment. Briefly, the mouse skeletal muscle tissue was processed exactly as described for FACS isolation of MuSCs up to the red blood cell lysis step. After red blood cell lysis, the remaining cells were centrifuged at 400 g for 15 min. The cell pellet was resuspended in 80 µL of FACS buffer and 20 µL of the Satellite Cell Isolation Kit (Miltenyi Biotec, #130-104-268). The cells were incubated for 15 min at 4 °C, after which they were passed through a MACS LS column (Miltenyi Biotec, #130-042-401) mounted on a QuadroMACS separator (Miltenyi Biotec, #130-091-051), and the flowthrough was collected. The column was washed twice with 1 mL FACS buffer, and this wash eluate was combined with the flowthrough of the cells, which was then centrifuged for 5 min at 400 g. The steps of labeling, magnetic depletion and collection of the flowthrough were repeated one more time with a fresh MACS LS column, to maximize the removal of non-MuSC cells. The flowthrough was centrifuged at 400 g for 5 min and resuspended in 80 µL of FACS buffer. 20 µL of anti-integrin α-7 microbeads (Miltenyi Biotec, #130-104-261) was added to the cell suspension and then incubated at 4 °C for 15 min, after which the mixture was passed through a fresh MACS LS column mounted on a QuadroMACS separator, to bind MuSCs to the column. The column was washed 3 times with 1 mL FACS buffer, and these eluates were discarded. After the washes, the LS column was removed from the magnetic separator and placed on top of a 15 mL collection tube. To collect MACS isolated MuSCs, 5 mL of FACS buffer was then applied to the LS column, and the magnetically labeled cells (i.e., α-7 integrin^+^) were flushed from the column into the collection tube using the plunger.

### MACS-and FACS-based Pax7^+^ cell enrichment purity assessment

To compare the purity of MuSCs isolated using FACS as compared to MACS, Pax7-nGFP mice were humanely euthanized, and their muscles were processed up to the point of incubation in 1 mL of red blood cell lysis buffer. After lysis, 10 mL of FACS buffer was added to the cell suspension, which was then evenly split into two 15 mL conical tubes. The tubes were each centrifuged at 400 g for 15 min. One of the tubes was processed based on the MACS isolation protocol, and the other was processed based on the FACS isolation protocol. The cells prepared by each method were then analyzed, using a FACS-Aria machine, with a pressure of 20Psi (low pressure) and 100 µm nozzle diameter, to determine the purity of the isolated cells, which was reported as the % total GFP^+^ (i.e., Pax7^+^) cells contained in each sample. Supplementary Figure 3 experiments that were conducted to determine the % total GFP^+^ cells enriched using a variety of optimisations in the MACS protocol were analyzed using the BD Accuri™ C6 Plus Flow Cytometer.

### Mini-MEndR assay

To conduct mini-MEndR assays, mini-myotube templates were fabricated and then cultured in MEndR DM for a period of 5 days, as described above. Towards the end of differentiation day 6, prospectively isolated MuSCs were resuspended in MuSC GM (Table 2) at a concentration of 212,500 cells/mL. Mini-myotube templates were then carefully removed from the 96-well plate using a sterile tweezer and gently placed, with their inverted side up, into the cavities of a sterile custom polydimethylsiloxane (PDMS) block. The PDMS block was placed within an ethanol-sterilized plastic container with a moist paper towel on the bottom to create a humidified chamber. 4 µL of resuspended MuSC solution (i.e., 850 cells per tissue) was dispensed onto the tissue surface, and evenly spread using a sterile cell spreader. The plastic container was then sealed with a tight-fitting lid and the stem cell laden myotube templates (i.e., mini-MEndR tissues) were then incubated for 1 hour at 37 °C, to allow the stem cells to incorporate into the mini-myotube template. Following the incubation, tissues were carefully transferred back into their respective 96-well plate positions containing MEndR DM supplemented with 5 ng/mL rh-FGF2, and either p38α/μ MAPKi or DMSO carrier control (Table 4). The media, containing MEndR DM, FGF2 and drug treatments, was completely replaced every other day over the next 5 days. For the remaining duration of the assay, tissues were cultured in MEndR DM, and half of the media was replaced every other day.

### Statistical Analysis

A minimum of three biological replicates (N) were conducted with multiple technical replicates (n) for most experiments as detailed in Table 5. Statistical analysis was performed using GraphPad Prism 8.0 software. For comparison of two variables, statistical differences were determined by unpaired Student’s t-test or unpaired t-test with Welch’s correction. For data with more than two variables compared, a one-way ANOVA followed by Tukey’s multiple comparison test was utilized (Table 5). All values are expressed as mean ± standard error of the mean (SEM). Significance was defined as p ≤ 0.05.

**Table 5.** Statistical tests and analysis.

| Figure | Biological experiments (N) | Total tissues/wells per condition (n) | Total images/wells per tissue | Statistical tests |
| --- | --- | --- | --- | --- |
| 1C | 1 per cell line | 3 | 5 | Ordinary one-way ANOVA with Tukey's multiple comparison's test |
| 1D | 1 per cell line | 3 | 5 | Ordinary one-way ANOVA with Tukey's multiple comparison's test |
| 1E | 1 per cell line | 3 | 5 | Ordinary one-way ANOVA with |
|  |  |  |  | Tukey's multiple comparison's test |
| 1H | 1 per cell line | 2 | 3 | Ordinary one-way ANOVA with Tukey's multiple comparison's test |
| 2C | 2 | 8 | 1 | Two-way ANOVA with Sidak's multiple comparisons test |
| 2F | Mixed in: 3<br>Pre-adsorped: 4 | Mixed in: 11<br>Pre-adsorped:24 | 1 | Unpaired two-tailed t-test |
| 3C | 3 | Day 7: 9<br>Day 10: 10<br>Day 14: 10 | 1 | Ordinary one-way ANOVA with Tukey's multiple comparison's test |
| 3D | 3 | Day 7: 9<br>Day 10: 8<br>Day 14: 6 | 1 | Ordinary one-way ANOVA with Tukey's multiple comparison's test |
| 4C | 6 | CTX (-): 18<br>CTX (+): 20 | 1 | Unpaired t-test |
| 4D | 5 | CTX (-): 15<br>CTX (+): 16 | 1 | Unpaired t-test |
| 5C | 5 | DMSO CTX (-):15<br>DMSO CTX (+):15<br>p38i CTX (+):15<br>EGFRi CTX (+):15 | 1 | Ordinary one-way ANOVA with Tukey's multiple comparison's test |
| 5D | 4 | DMSO CTX (-):12<br>DMSO CTX (+):12<br>p38i CTX (+):12<br>EGFRi CTX (+):10 | 1 | Ordinary one-way ANOVA with Tukey's multiple comparison's test |
| 6C | 4 | DPI7 DMSO: 11<br>DPI7 p38i:12<br>DPI10 DMSO:11<br>DPI10 p38i:10 | 1 | Multiple t-tests with Welch's correction |
| 6D | 3 | DPI7 DMSO: 9<br>DPI7 p38i:9 | 1 | Multiple t-tests with Welch's correction |
|  |  | DPI10 DMSO:8<br>DPI10 p38i:9 |  |  |
| 7C | 3 | DMSO: 6<br>p38i: 8 | 4-7 | Unpaired two-tailed<br>T-test with Welch's<br>correction |
| 7D | 3 | DMSO: 7<br>P38i: 8 | 4-7 | Unpaired two-tailed<br>T-test with Welch's<br>correction |
| 7E | 3 | DMSO: 6<br>p38i: 8 | 4-7 | Unpaired two-tailed<br>T-test with Welch's<br>correction |
| 7F | 3 | DMSO: 7<br>P38i: 8 | 4-7 | Unpaired two-tailed<br>T-test |
| SI 4B | 3 | FACS: 9<br>MACS: 10 | 1 | Unpaired two-tailed<br>T-test with Welch's<br>correction |

## Results

### Selection of a commercially available cell source for myotube template fabrication

As a first step in establishing a workflow that removed bottlenecks in manufacturing observed in the original MEndR protocol^10^, we set out to select a commercially available primary human myoblast cell line to remove logistical challenges associated with isolating cells from fresh patient tissue samples. To do this, we differentiated three commercially available primary human myoblast cell lines (Table 1) in 2D culture and compared them across a battery of metrics to select lines for our study that were maximized for differentiation potential and we expected would be minimized for Pax7^+^ “reserve cell”-mediated self-repair^10,16^. Specifically, we plated primary myoblasts at equivalent passage number and cultured them for 1 day under growth media (GM) conditions. We then transitioned the cultures to differentiation media (DM), and on the 9th day of differentiation, we fixed the cultures for analysis (Figure 1A). All three cell lines formed multinucleated structures that were immunoreactive for the contractile apparatus protein, sarcomeric α-actinin (SAA; Figure 1B). The 18M cell line produced SAA^+^ structures that covered a greater proportion of the culture surface area as compared to the other two lines (Figure 1C). We found that all three cell lines performed equivalently with regards to myotube nuclear accretion upon evaluating the proportion of nuclei that contributed to SAA^+^ structures (Figure 1D). These data, coupled with visual inspection of the SAA staining (Figure 1B), in turn suggested that the 18M cell line likely produces myotubes of greater diameter than those produced by the other two lines. Using immunocytochemistry to visualize Pax7^+^ cells, we found that both the 18F and 19F cell lines had a greater proportion of mononucleated Pax7^+^ ‘reserve cells’^16^ present in myotube cultures when compared to the 18M line (Figure 1E). Finally, we confirmed that the cell lines were competent to produce myotubes when resuspended in a 3D fibrin-based hydrogel and cultured in a cellulose scaffold using the original myotube template manufacturing protocol (Figure 1F-H). In all cases, we observed the formation of myotubes structures and similar levels of SAA-coverage within the scaffold (Figure 1G-H). Since our prior studies suggested that high reserve cell content can mask the regenerative influence of muscle stem cells added to the assay^10^, we elected to move forward with the 18M and 19F lines and abandon the 18F line for this study.

**Figure 1.**
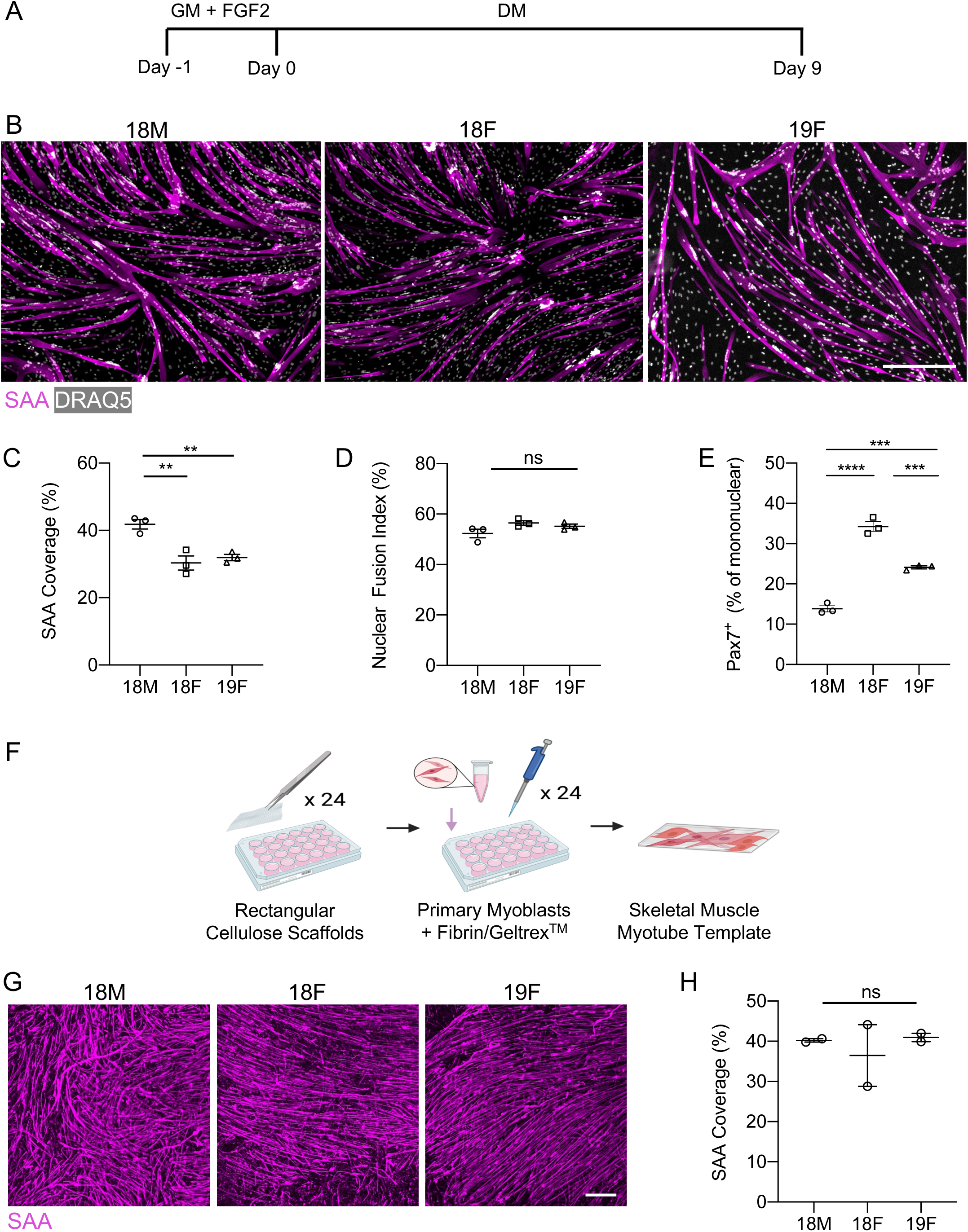
Characterization of commercially available human pMB lines. **(A)** Experimental timeline for the 2D differentiation of the human primary myoblasts. **(B)** Representative 10X confocal images of the three commercially available human myoblasts immunostained for sarcomeric α-actinin (SAA; magenta) and counterstained for DRAQ5 (grey) on day 9 of differentiation in 2D culture. Scale bar, 500 µm. **(C-E)** Bar graphs quantifying SAA coverage **(C**), average nuclear fusion index **(D)**, and ratio of Pax7^+^ reserve cells amongst the mononuclear cells **(E)** in 2D culture on day 9 differentiation comparing the three cell lines. Graphs display mean ± s.e.m., one-way ANOVA with Tukey post-test. ** p < 0.01, *** p < 0.001, **** p < 0.0001, ns. - no significance. n=3 wells from N=1 experiment per cell line. **(F)** Schematic of the 24-well myotube template derivation workflow. **(G)** Representative confocal images of SAA (magenta) immunostaining in myotube templates at day 8 of differentiation that were generated using the 18M, 18F and 19F cell lines. Scale bar, 500 µm. **(H)** Bar graph quantification of myotube template SAA coverage at day 8 of differentiation. Graphs display mean ± s.e.m., one-way ANOVA with Tukey post-test, ns. – no significance. n=2 tissues from N =1 experiment per cell line.

### Cellulose scaffolds pre-adsorbed with thrombin to allow scalable manufacturing

Having selected commercial cell lines, we set out to establish a workflow for production of myotube templates in a manner that mitigated time-sensitive steps in the original fabrication protocol that we have observed prove challenging for new users that want to adopt the MEndR platform. In the original protocol, myogenic progenitors were resuspended within a fibrinogen / thrombin / Geltrex^TM^ based hydrogel precursor solution, which was subsequently manually distributed over the cellulose scaffold to evenly spread the hydrogel suspensions and encourage infiltration into the scaffold pores prior to hydrogel gelation^10^. A fibrin clot (i.e., hydrogel) forms rapidly upon combining the fibrinogen with thrombin, allowing only a short window of time to resuspend the myogenic progenitors within the solution, dispense the gel-cell solution onto the scaffold, and infiltrate into the cellulose pores. When fabricating only a small (<24) number of scaffolds, this time window is a minimal problem. However, we were concerned that when fabricating 96-scaffolds, this short time window would present a significant challenge. To decouple the hydrogel mixing and gelation steps, we assessed if we could adsorb thrombin (which initiates fibrin gelation), to the cellulose scaffold (Figure 2A). We reasoned that using this approach the myogenic progenitor / fibrinogen / Geltrex^TM^ solution would polymerize in situ only upon making contact with the thrombin-adsorbed cellulose scaffold. We first wanted to confirm that pre-adsorbing fibrin on the scaffold resulted in hydrogel gelation upon infiltration into the scaffold. To this end, we mixed fluorescent beads into the fibrinogen / thrombin / Geltrex^TM^ hydrogel precursor solution, infiltrated the solution into the pre-adsorbed scaffold and allowed gelation, imaged the beads, washed vigorously to rinse out any unpolymerized hydrogel (and beads), and then re-imaged the remaining beads to allow us to quantify bead loss due to the washing step (Figure 2A-C). As expected, when no thrombin was present, bead loss occurred upon washing the scaffolds. In conditions where thrombin was directly mixed into the pre-cursor solution, or pre-adsorbed onto the cellulose scaffold, minimal bead loss was observed (Figure 2B-C).

**Figure 2.**
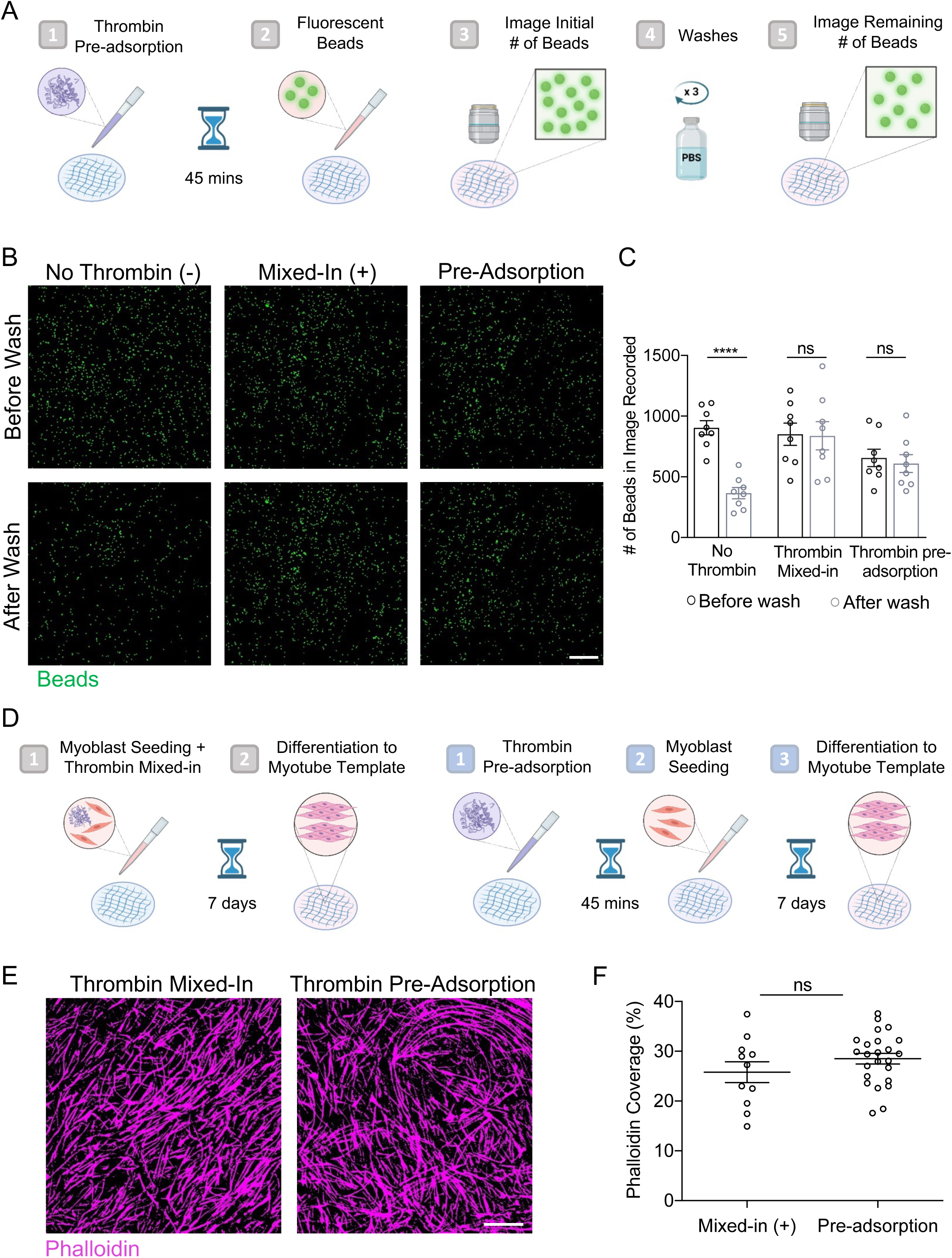
Preabsorbing thrombin in paper scaffolds to facilitate myotube template seeding. **(A)** Schematic of the bead assay workflow. Cellulose scaffolds were pre-adsorbed with thrombin and seeded with fluorescent beads diluted 10 times in ECM master mix. The beads were imaged before and after washing to quantify bead retention. **(B)** Representative confocal images of bead assay paper scaffolds before and after washing. Scale bar, 500 µm. **(C)** Quantification of the number of beads before washing (black) and after washing (grey) for fluorescent beads seeded on paper scaffolds without thrombin, with 0.8 U/ml thrombin mixed-in, or with 0.8 U/ml thrombin pre-adsorption. Graphs display mean ± s.e.m., two-way ANOVA followed by Sidak’s multiple comparisons test *** p < 0.001, N=2, n=8. **(D)** Experimental workflow for comparing thrombin mixed-in (left) to pre-adsorption (right) methods on myotube template formation. **(E)** Representative confocal images of myotube template stained for phalloidin (magenta) after 7 days of differentiation. Scale bar, 500 µm. **(F)** Quantification of phalloidin coverage for thrombin mixed-in vs pre-adsorption methods. ns. = no significance. n=11-24 tissues from N=3-4 independent experiments.

When we used this pre-adsorption protocol with myoblasts rather than beads, within one week of culture, we observed thin sheets of multinucleated myotubes form across the two cell lines tested (Figure 2D-E). Further, the proportion of 3D tissues covered by SAA^+^ myotubes in myotube templates formed using pre-adsorption of thrombin to the scaffold was similar to those formed when thrombin was directly mixed in (Figure 2E-F). We therefore conclude that the pre-adsorption of the thrombin in the cellulose scaffold enabled robust manufacturing of muscle templates that differentiated as expected.

### Fabrication of myotube templates within a 96-well footprint

We next set out to manufacture MEndR tissues compatible with a 96-well plate footprint. To miniaturize myotube templates from the original 24-well rectangular format^10^, we cut sheets of cellulose teabag paper into circular discs using a 5 mm circular biopsy punch (Figure 3A). The circular cellulose scaffolds were autoclaved and then placed into pluronic acid treated wells of a 96-well plate using anti-static tweezers. The pluronic acid treatment discouraged myoblast migration or attachment beyond the boundary of the scaffold (Supplementary Figure 1). For both of the myoblast cell lines, we observed formation of multinucleated myotubes by day 7 of differentiation and similar levels of SAA tissue coverage at days 10 and 14 of culture, at which point, we concluded the experiment (Figure 3B-D). SAA tissue coverage at day 7 in the miniaturized format (Figure 3C-D) was similar to that observed in the large format at day 8 for both cell lines (Figure 1H versus Figures 3C and D). Importantly, we found that, in general, three rounds of practice seeding the 96-well format myotube templates using the optimized protocol was sufficient for new users to master the method (Supplementary Figure 2). We therefore concluded that myotube templates could be generated in this smaller dimension format and that new users with myoblast cell culture experience can quickly become proficient with the method.

**Figure 3.**
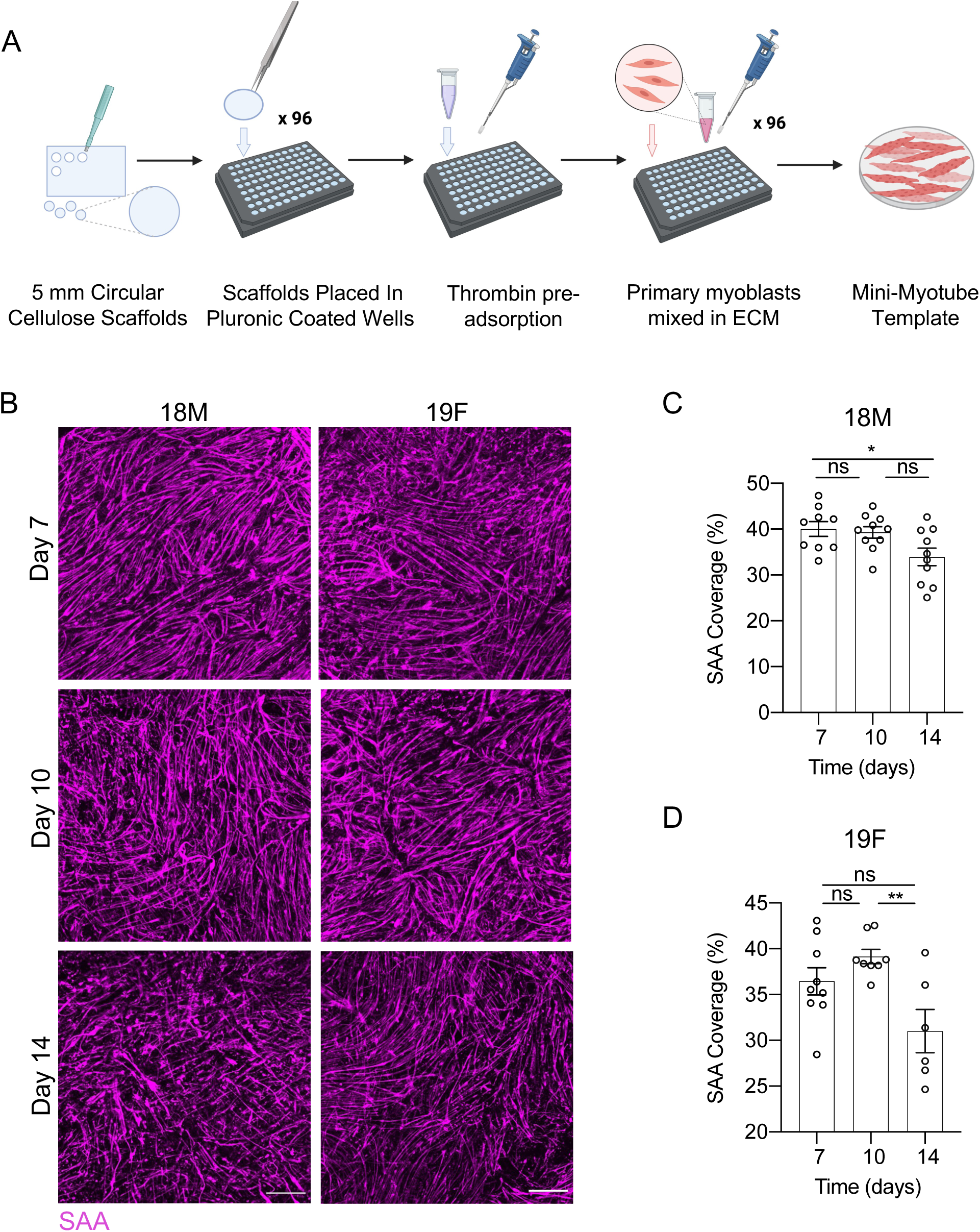
Mini-myotube template derivation. **(A)** Schematic of the mini-myotube template derivation workflow. Cellulose scaffolds were pre-adsorbed with thrombin and seeded with human pMBs mixed in an ECM mixture containing DMEM, fibrinogen and Geltrex^TM^. **(B)** Representative confocal images of SAA (magenta) immunostaining for mini-myotube templates generated with the 18M and 19F cell lines and analyzed on days 7, 10, and 14 of differentiation. Scale bar, 500 µm. **(C, D)** Quantification of SAA coverage in mini-myotube templates generated with the 18M **(C)** and 19F **(D)** cell lines on days 7, 10, and 14 of differentiation. Graphs display mean ± s.e.m. one-way ANOVA with Tukey post-test. * p < 0.05, ** p < 0.01 ns. = no significance. n = 6-10 tissues from N = 3-4 independent experiments per cell line.

### Miniaturized myotube templates possess limited self-repair capacity

Having miniaturized our myotube template platform into a 96-well format, we next set out to demonstrate that our miniaturized platform supported the stem cell mediated repair workflow. Specifically, we wanted to confirm that the miniaturized myotube templates were susceptible to damage induced by cardiotoxin, a snake venom toxin commonly used to induce muscle damage in mouse studies^17^. To do this, on day 7 of mini-myotube template culture, we introduced cardiotoxin to the culture media (Figure 4A). Mini-myotube templates were fixed 24 hrs after the initial cardiotoxin exposure and immunostained to visualize and quantify the resulting SAA^+^ myotube structures (Figure 4A-D). We observed a 21.31% +/-4.37% and 24.12% +/-5.24% decrease in SAA immunoreactivity one day after cardiotoxin treatment in myotube templates generated from the 18M and 19F myoblast lines, respectively (Figure 4C-D), consistent with prior results obtained using the larger myotube template format^10^.

**Figure 4.**
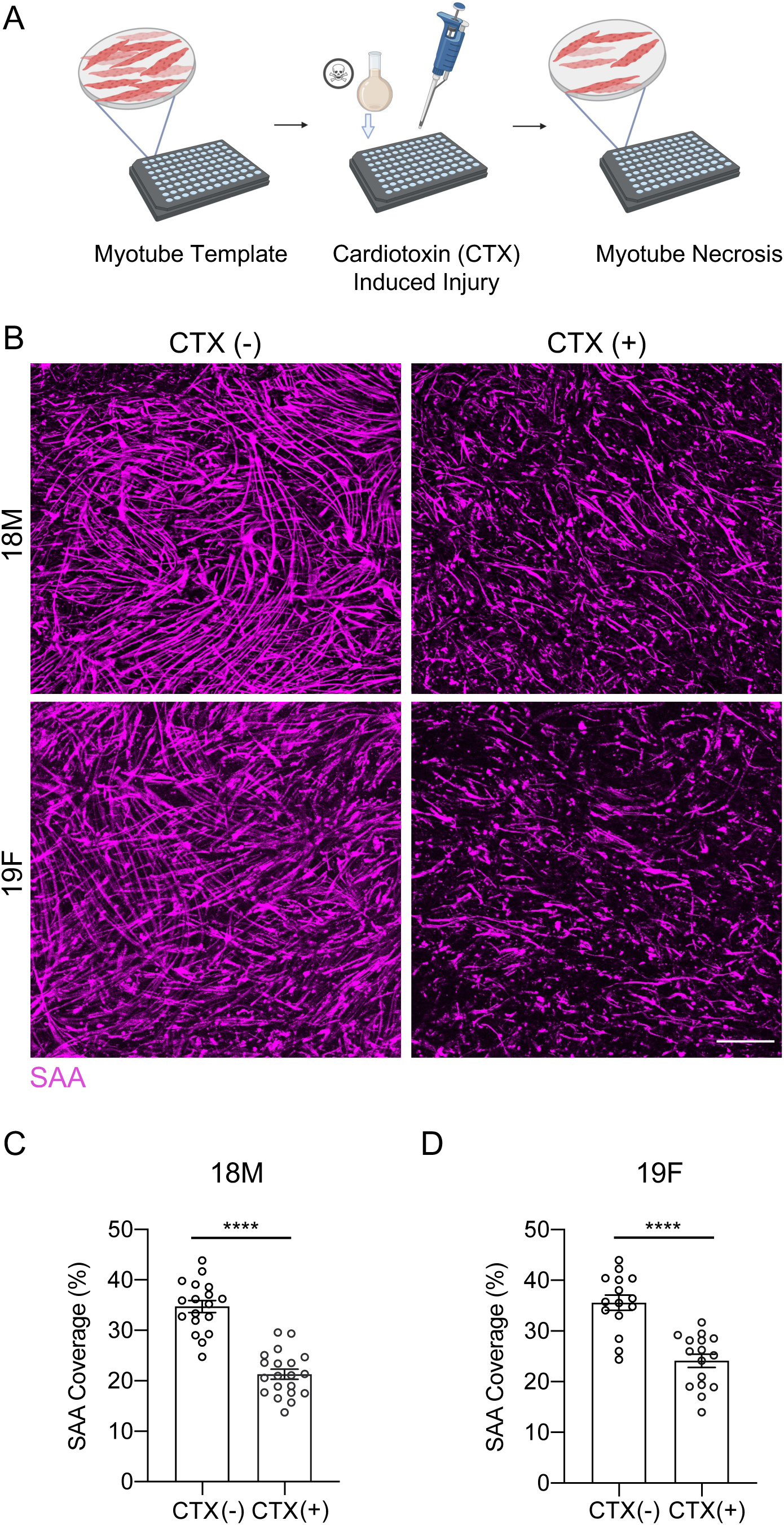
Myotoxin-induced mini-myotube templates injury. **(A)** Schematic of the template injury workflow. Mini-myotube templates are incubated in MEndR differentiation media containing 0.5 μm cardiotoxin (CTX) for 4 hr at 37 °C on an orbital shaker. **(B)** Representative confocal images of uninjured (left; (-)) and injured (right; (+)) mini-myotube templates generated with the 18M (top) or 19F (bottom) cell lines (SAA; magenta) at 1 DPI. Scale bar, 500 µm. **(C, D)** Quantification of SAA tissue coverage in uninjured and injured mini-myotube templates generated with 18M **(B)** or 19F **(C)** cell lines at 1 DPI. Graphs display mean ± s.e.m. Unpaired student’s t-test, **** p < 0.0001. n =11-20 tissues from N=5-6 independent experiments per cell line.

We next analyzed the mini-myotube templates 10 days after cardiotoxin exposure, to determine whether Pax7^+^ reserve cell activity alone regenerated the injured myotube templates (Figure 5A). Similar to the larger myotube template format^10^, we found that the SAA content failed to return to levels observed in the uninjured control (Figures 5B-D, DMSO CTX(-) vs CTX (+)). As expected, when templates were treated with an inhibitor of p38 α/β MAP kinase (p38i), a treatment known to expand both muscle stem and progenitor cells^8^, we observed that SAA content was able to return to levels matching the uninjured control (Figure 5B-D). By contrast, treatment with an epidermal growth factor receptor inhibitor (EGFRi) failed to stimulate reserve cell-mediated myotube template repair, as measured by SAA coverage (Figure 5B-D). Together, these results confirmed that the mini-myotube templates were receptive to injury, and had limited self-repair capacity, allowing for enhancement by engrafting freshly isolated MuSCs.

**Figure 5.**
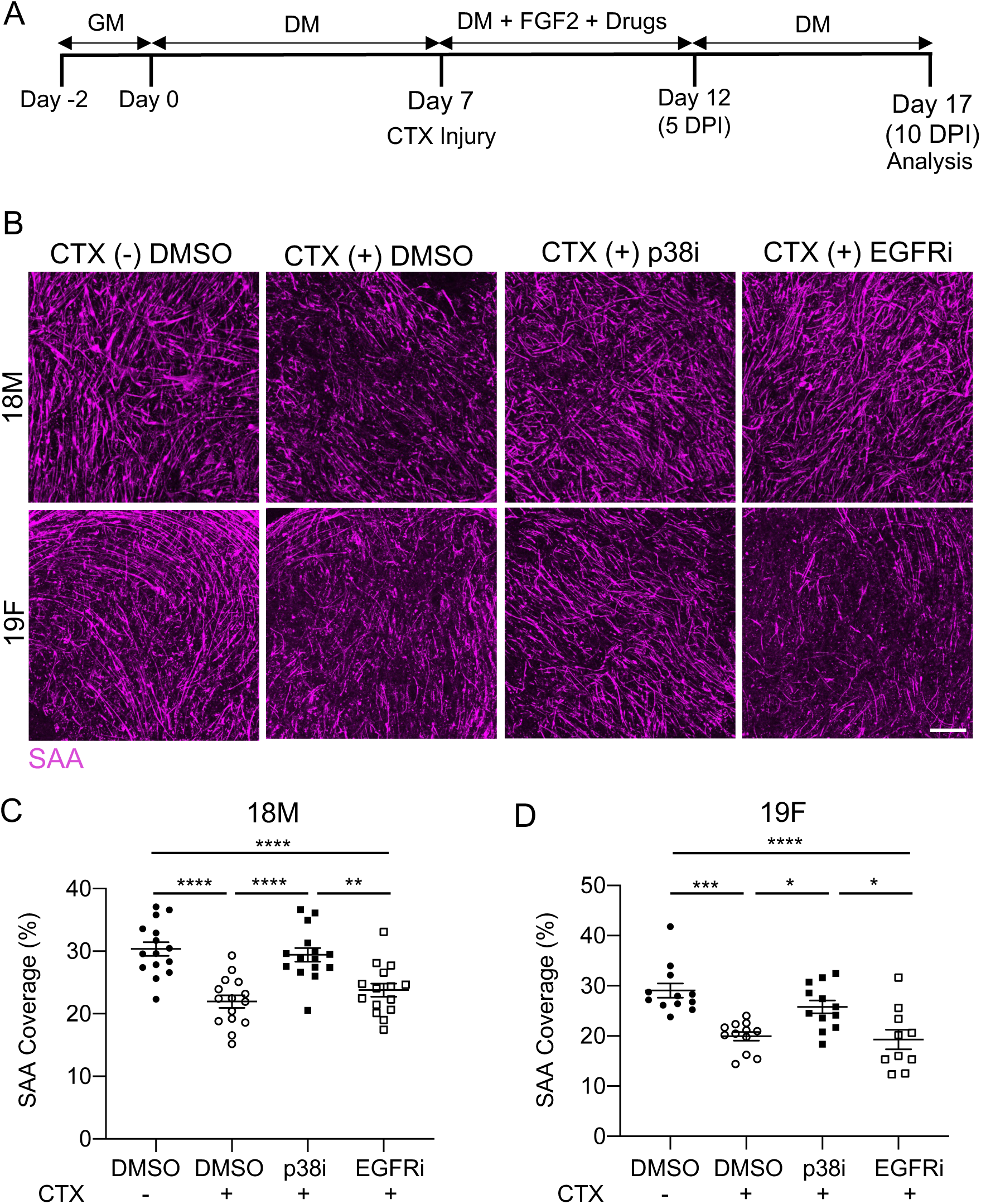
p38α/β MAPKi, but not EGFRi, induces mini-myotube template self-repair. **(A)** Experimental timeline to study mini-myotube template self-repair capacity. **(B)** Representative confocal images of uninjured (CTX(-)) and injured (CTX(+)) myotube templates generated with 18M (top) or 19F (bottom) cell lines in response to treatment with p38α/β MAPK inhibitor (p38i), epidermal growth factor inhibitor (EGFRi) or DMSO carrier control at 10 DPI. Scale bar, 500 µm. (**C, D)** Quantification of SAA coverage in uninjured and injured myotube templates generated with 18M **(C)** or 19F **(D)**. Graphs display mean ± s.e.m. one-way ANOVA with Tukey post-test, *p<0.05, ** p < 0.01, *** p < 0.001, **** p < 0.0001. n =10-15 tissues from N=4-5 independent experiments per cell line.

### A MACS-based MuSC enrichment strategy supports MEndR assay scale-up

We next set out to demonstrate that muscle stem cell mediated skeletal muscle tissue repair occurs in the miniaturized myotube template format. We previously showed that freshly FACS-sorted MuSCs introduced to injured myotube templates contribute to the process of tissue repair by producing new myotubes^10^. For these studies, we prospectively isolated MuSCs from enzymatically digested murine skeletal muscle tissue using a magnetic-activated cell sorting (MACS) strategy (Supplementary Figure 3), which we first confirmed to produce similar results as compared to MuSCs isolated using a traditional fluorescence activated cell sorting (FACS) protocol (Supplementary Figure 4). For this comparison, we harvested hindlimb skeletal muscle tissue from humanely euthanized Pax7-nGFP transgenic reporter mice (Supplementary Figure 3) or those ubiquitously expressing yellow fluorescent protein (YFP; Supplementary Figure 4) and subjected it to standard mechanical dissociation and enzymatic digestion protocols to produce a single cell suspension. The cell suspension was split equally, processed, and sorted (Supplementary Figure 3A) using either a FACS (Supplementary Figure 3B-C) or MACS-based protocol (Supplementary Figure 3C-D). The cell populations harvested from Pax7-nGFP animals were characterized immediately after enrichment using flow cytometry (Supplementary Figure 3C). We found that 97.87% +/-2.24% of events from FACS-enriched events were GFP^+^, whereas 91.70% +/-0.97% of events were GFP^+^ following a modified MACS enrichment strategy that employed two-rounds of lineage depletion (Supplementary Figure 3C-D). We speculated that the greater proportion of GFP-negative cells within the MACS-enriched cell population could be owed to integrin α-7^+^ contaminating cell types^18^, or that it may be an artifact of magnetic-enrichment protocols which are less effective than FACS at removing cellular debris. In prior studies, we plated the FACS and MACS-enriched cell populations in culture and then 12 hrs later, enumerated the viable GFP positive and negative cells in each well^19^. This uncovered a dramatic loss of the GFP-negative population from the MACS-enriched cell population, resulting in an enrichment for GFP-positive cells that matched values obtained using FACS-enrichment. As expected, increased experience using the MACS method correlates with a consistently higher proportion of GFP^+^ cells in the populations enriched from Pax7-nGFP transgenic animals (data not shown). It is also worth noting that in our experience MACS-based enrichment consistently offered higher cell yields when compared to the FACS-based strategy (data not shown). Together this data suggested that MACS was a feasible and convenient method for sorting MuSCs from mouse hindlimb skeletal muscle tissue for our scaled MEndR assay.

### Miniaturized format supports muscle endogenous repair (mini-MEndR) assays

We next wanted to confirm that our miniaturized myotube template system, seeded with MuSCs, could replicate key results observed using the original MEndR assay which successfully identified and recapitulated outcomes of known stimulators of murine muscle stem cell mediated muscle tissue repair^10^. We generated myotube templates, and then on day 6 of differentiation we introduced MACS enriched MuSCs from YFP-transgenic animals. 12 hr after MuSC engraftment, co-cultures were then exposed to cardiotoxin to injure the myotube templates, after which we introduced to the culture media a p38 α/μ MAPK inhibitor (p38i), a known modulator of stem cell skeletal mediated repair, or DMSO carrier control. At 7 or 10 days post-injury (DPI), the tissues were fixed and immunostained to evaluate the donor-derived contributions to muscle tissue repair (Figure 6A-B). Consistent with our results in the large MEndR format, we observed increased donor derived myotube formation in tissues treated with p38i, suggesting enhanced stem cell mediated repair that was apparent at 10 DPI (Figure 6C-D). This effect was observed when MuSCs were introduced to myotube templates generated from both the 18M and 19F primary myoblasts (Figure 6C-D). Further, MuSCs prospectively isolated using the MACS-enrichment method performed in a functionally equivalent manner by this metric when compared against those sorted using a classic FACS-based method (Supplementary Figure 4). The effect of p38i on donor myotube production was statistically detectable by 7 DPI, and was not significantly different from the results obtained at 10 DPI. This observation suggests that a shortened timeline to assess stem cell-mediated muscle production using the mini-MEndR assay is feasible, benefiting future screening applications.

**Figure 6.**
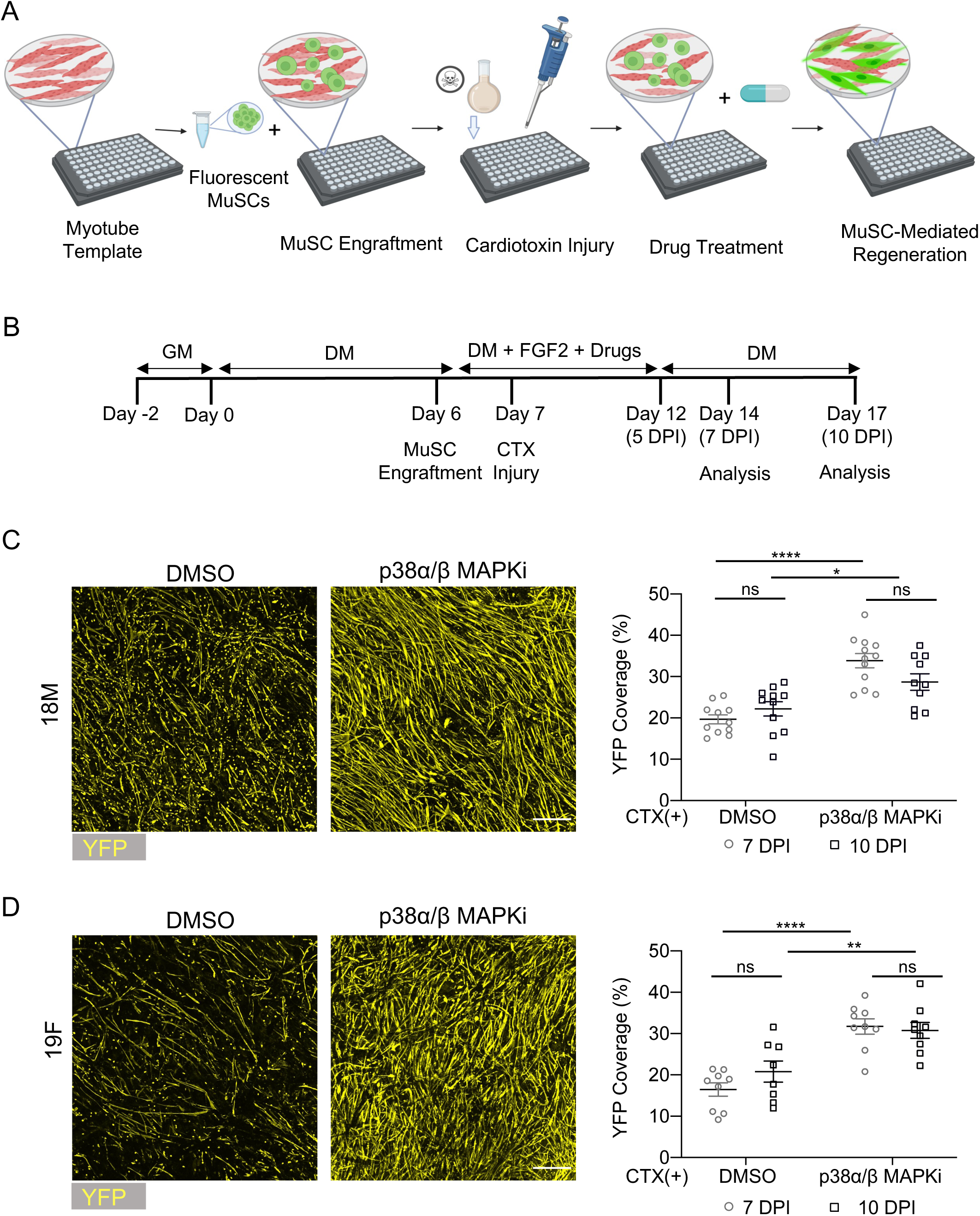
p38α/β MAPKi treatment augments MuSC-mediated muscle production in mini-MEndR assay. **(A)** Schematic of the mini-MEndR assay workflow. MuSCs from YFP transgenic mice are isolated using the adapted magnetic assisted cell sorting (MACS) protocol and then engrafted onto the mini-myotube templates on day 6 of differentiation, followed by drug treatments and cardiotoxin-induced tissue injury. **(B)** Detailed experimental timeline for the mini-MEndR assay workflow. **(C)** Representative confocal images of MuSC-mediated (YFP**^+^**) myotube production in the injured mini-myotube templates produced with 18M (top panel) 19F (bottom panel) in response to treatment with DMSO or p38α/β MAPKi harvested at 7 DPI. Scale bar, 500 µm. (**D-E)** Quantification of the MuSC-mediated regeneration (%YFP coverage) in the injured mini-myotube templates generated with 18M **(D)** or 19F **(E)** cell line harvested at 7 DPI or 10 DPI. Graphs display mean± s.e.m; Unpaired two-tailed T-test with Welch’s correction was conducted for each condition. * p<0.05, ** p < 0.001, **** p, n=9-12 tissues from N=3-4 independent experiments per cell line.

We next assessed another key read-out of stem cell mediated skeletal muscle repair, niche repopulation, which we have shown is captured in the original MEndR format^10^. For these studies, we repeated the protocol described above, using instead MuSCs isolated from Pax7-nGFP transgenic reporter animals (Figure 7A). On day 7 post-injury, when tissue repair was complete, we observed a population of mononucleated Pax7^+^ donor-derived (GFP; Figure 7B), which we previously showed had reversibly exited cell cycle^10^. Further, and in line with results obtained in the original MEndR platform, we observed a ∼3-fold increase in these donor-derived Pax7^+^ cells in response to p38i treatment (Figure7C-D). The MEndR and mini-MEndR assays were designed as a mixed species cell ‘competition’ assay in part to enable multiplexed analyses of human and mouse Pax7^+^ populations within the same assay. The mouse Pax7^+^ cell population is isolated from a transgenic animal and therefore easily distinguished from the unlabeled Pax7^+^ ‘reserve cells’ that exist within the primary human myoblast population, which is used to fabricate the myotube template. Thus, enumerating the mononucleated Pax7 immunoreactive cells present in the myotube template at the end-point of a mini-MEndR assay, while tracking whether they are GFP labeled (mouse) or unlabeled (human) enables a cursory assessment of species specific Pax7^+^ cell responses. Consistently, when we enumerated the Pax7^+^ reserve cell abundance at the assay end-point, treatment with the p38i produced an increase in the incidence of mononucleated Pax7^+^ reserve cells (Figure 7E-F).

**Figure 7.**
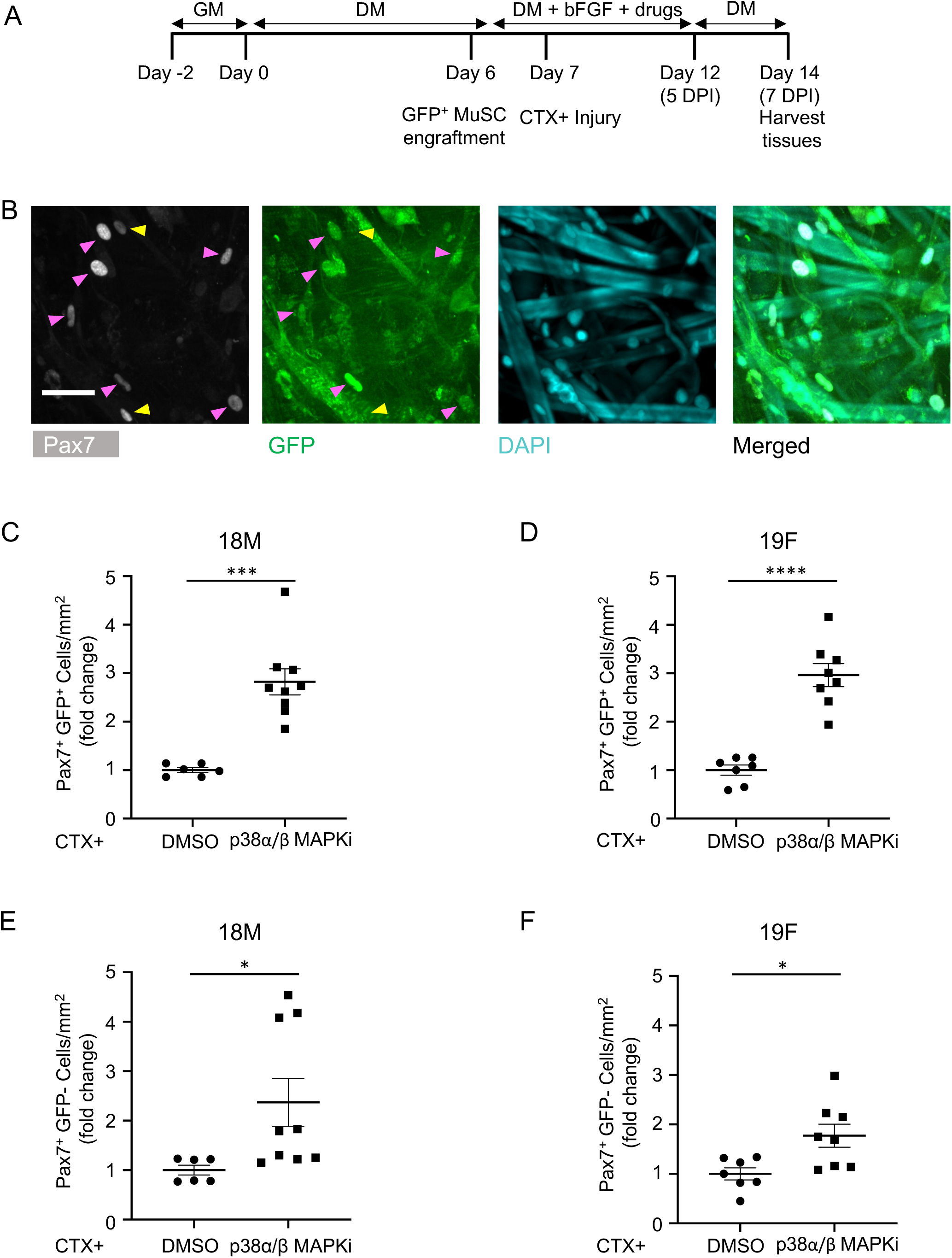
mini-MEndR assay captures response of treatment known to stimulate MuSC self-renewal. **(A)** Detailed experimental timeline of donor-derived MuSC niche repopulation. **(B)** Representative confocal images of Pax7^+^GFP^+^ donor derived MuSCs in mini-MEndR tissues treated with p38α/μ MAPKi at 7 DPI. Tissues are immunostained for Pax7 (grey), GFP (green), and DAPI (cyan). Magenta arrowheads point to the Pax7^+^ GFP^+^ donor MuSCs and yellow arrowheads point to the Pax7^+^ GFP^-^ reserve cells derived from the myotube template. Scale bar, 50 µm. **(C-F)** Quantification of Pax7^+^ GFP^+^ cells derived from donor MuSCs (**C-D**) or Pax7^+^ GFP^-^ reserve cells derived from the myotube template (**E-F**) in mini-MEndR tissues generated with 18M **(C, E)** and 19F **(D, F)** cell lines, following treatment with DMSO or p38α/β MAPKi in the context of injury and analyzed at 7 DPI. Graphs display mean ± s.e.m.; Unpaired two-tailed T-test with Welch’s correction. * p < 0.5, *** p < 0.001, **** p < 0.0001. n=6-9 tissues from N=3 independent experiments per cell line.

Together this data suggested that the miniaturized muscle endogenous repair assay (mini-MEndR) replicates all of the stem cell functional metrics previously reported in the original large format MEndR assay.

## Discussion

In this paper we set out to miniaturize and improve the manufacturing workflow of our previously reported skeletal muscle stem cell mediated repair assay, known as MEndR^10^, to enable wider adoption of this powerful assay to identify novel molecular targets that promote muscle regeneration. We addressed three critical bottlenecks in the workflow of the original MEndR assay, that have up until now limited its capacity to assess molecular targets at a practically useful scale. Firstly, we demonstrate use of primary human myoblasts from a commercial source to generate the myotube template to reduce myotube template variation associated with the use of different muscle biopsy donors. Second, we developed a method to integrate the hydrogel crosslinker into the scaffold to trigger in situ gelation to counter the rapid gelation properties of the fibrin hydrogel used to manufacture the myotube template, which had limited the number of myotube templates that can be robustly manufactured using one batch of myoblast-hydrogel suspension. Third, we validate the use of an alternative to FACS for sorting skeletal muscle stem cells from digested skeletal muscle which overcomes limits on the number of wells that can be assessed in parallel using freshly isolated adult stem cells. Further, we miniatured the format of the assay ∼6-fold to reduce both the number of template cells and added stem cells required per well to perform the assay, and demonstrated the possibility of a shorter assay endpoint.

The use of cross-linker adsorption to the paper scaffold to enable in situ hydrogel gelation during tissue manufacturing was particularly critical to able scalable manufacturing of the myotube template due to the rapid gelation of fibrin upon addition of the thrombin crosslinker to the bulk hydrogel solution containing the myoblasts. We anticipate this innovation could enable the manufacturing of myotube templates using robotic liquid handling in the future to support tissue manufacturing for large scale screening efforts. Indeed this strategy could potentially be utilized to generate tissues for other applications that use two component hydrogels in which gel crosslinking is initiated by the addition of an external crosslinking component as opposed to simply a temperature change alone. Interestingly, we observed that the use of thrombin pre-adsorption dramatically reduced the number of attempts required to master the mini-MEndR seeding protocol, which suggests that pre-mature hydrogel gelation could be a key source of inter-well variation in other paper-based systems^13,20^.

While this work successfully increased the throughput capacity of our MEndR assay to enable potential screening of drug candidates at a 96-well plate scale, it is important to note some remaining limitations of the mini-MEndR assay. We note that removal of the myotube template from each well to integrate the muscle stem cells remains a rate limiting step in our work flow. However, this step does not significantly impact usability at a 96-well plate throughput. A 96-well throughput is appropriate for assessing 10s to 100s of targets. This manual removal step, however would limit use of robots in fully automating our assay. If a larger scale screening capacity is desirable in the future, the MEndR assay could be further integrated with the SPOT platform^20^, reported by our group, which we recently demonstrated is compatible with robotic liquid handler to enable massive increases in scale^21^.

Given that the myotube template contains a population of Pax7^+^ reserve cells that have the potential to contribute to myotube template repair, even in the absence of adding a MuSC population, means that readouts of mini-MEndR SAA-coverage alone are not able to distinguish between drugs that target this reserve cell population versus those that target the MuSC population specifically. Thus, it is important for MuSCs that are added to mini-MEndR experiments to be fluorescently labeled to disentangle MuSC and template reserve cell regenerative activities. An advantage of conducting mixed species mini-MEndR experiments is the ability to track the influence of treatments on both human Pax7^+^ reserve cell and mouse MuSC self-renewal and differentiation within the same experiment. The user has complete control over MuSC content in each experiment, but limited control over the drift in Pax7^+^ reserve cell content. Further, there is ongoing debate with regards to drawing conclusions about human MuSC biology based solely on human Pax7^+^ reserve cell studies. For example, the EGFRi treatment induced different effects on mouse MuSCs as compared to human Pax7^+^ reserve cells based upon our experiments, and whether this reflects an influence of species or is a MuSC vs reserve cell difference has yet to be determined. Thus, downstream studies to evaluate the effects of ‘hits’ on bonefide human MuSCs are to be expected, and adaptions of the assay to integrate MuSC freshly enriched from human muscle biopsies are underway. The capacity to conduct follow up experiments at a higher scale in combination with the compatibility of mini-MEndR tissues to be imaged semi-automatically with high-content imaging systems will serve to facilitate follow up dose response and mechanistic studies of molecular targets identified in the initial screen.

## Conclusions

Identification of drugs to boost the capacity of MuSCs to drive skeletal muscle endogenous repair is an exciting approach to address various conditions that lead to impaired skeletal muscle regeneration, such as aging or DMD. The MEndR assay provides a powerful functional assay “in a dish” with the potential to shortlist molecular targets that enhance human skeletal muscle endogenous repair. Here we adapted the original MEndR manufacturing workflow to enable the robust manufacturing of a 96-well plate “mini-MEndR” platform. Further we validated that this miniaturized platform recapitulates the known endogenous repair readouts captured in the original larger MEndR format. The mini-MEndR platform has the potential to enable the identification of novel molecular targets to modulate stem-cell mediated skeletal muscle repair and to facilitate an exploration of their cellular and molecular mechanism(s) of action.

## Acknowledgements

N.G., S.D., and B.X. contributed equally to this work. S.R., and E.J. contributed equally to this work. BioRender was utilized to produce schemata elements. This project was funded by a Canada First Research Excellence Fund “Medicine by Design (MbD)” grant to P.M.G. and A.P.M. (MbDC2-2019-02), a Natural Sciences and Engineering Research Council of Canada (NSERC) Idea to Innovation grant awarded to A.P.M. and P.M.G. (I2IPJ 549768-20), a University of Toronto Connaught Fund awarded to A.P.M. and P.M.G., a Canada Research Chair in Endogenous Repair award to P.M.G. (#950-231201), Stem Cell Network grant to P.M.G. (#FY20/DRP-1), Ontario Institute for Regenerative Medicine grant to P.M.G. (#2018-0510), CGS-D Awards to B.X. and E.J., a CGS-M Award to S.R., an NSERC CREATE TOeP scholarship to S.D., Ontario Graduate Scholarships to B.X., S.R., and E.J. and a Norman F. Moody Award to B.X.

## Author contributions

N.G., S.D., B.X., S.R., E.J., J.P., and A.F. designed and performed research, analyzed data, and prepared figures. A.P.M., P.M.G., and S.D. conceived of the project and supervised the research. All authors contributed to data interpretation. A.P.M., P.M.G., N.G., B.X., and S.R. wrote the manuscript. All authors reviewed and approved the manuscript.

## Conflict of Interest

The authors have no competing interests, or other interests that might be perceived to influence the results and/or discussion reported in this paper.

## Data and Materials Availability Statement

The datasets generated during and analyzed in this study are available from the corresponding author on reasonable request.

---

**Supplementary Figure 1.**
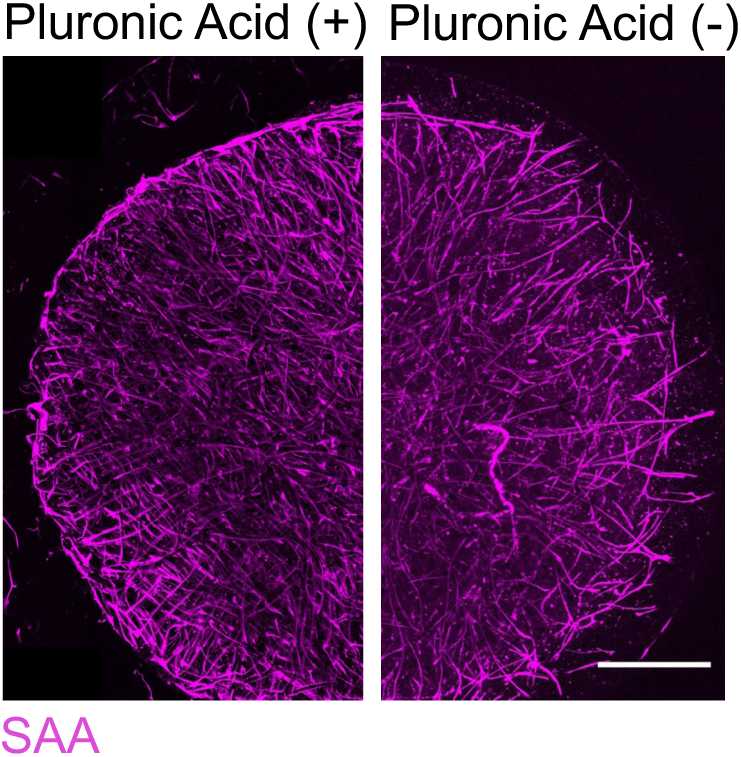
Pluronic acid coating allows for the generation of smooth mini-myotube template edges. Representative confocal images of mini-myotubes generated with (+) and without (-) pluronic acid coating. Tissues are immunostained for sacromeric α-actinin (SAA; magenta). Scale bar, 1 mm.

**Supplementary Figure 2.**
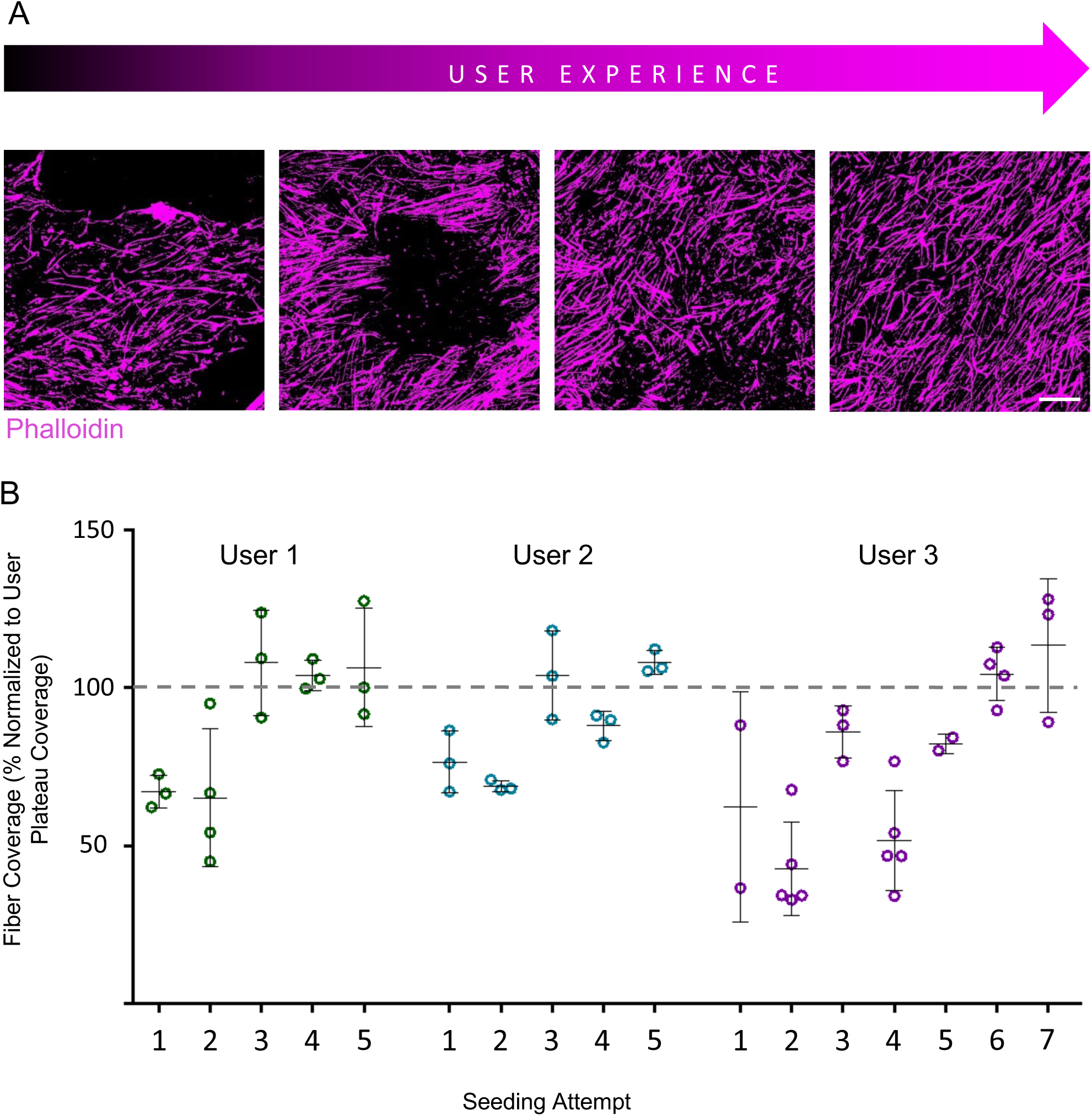
Mini-MEndR platform manufacture can be quickly mastered by new users. **(A)** Representative confocal images of myotube templates generated by the same user at different levels of experience. Tissues were fixed after 7 days of differentiation and stained with phalloidin (magenta). Scale bar, 500 μm. **(B)** Quantification of Phalloidin (User 1) or SAA (User 2, 3) tissue coverage attained by three different users who trained from first-time user to expert level as indicated by plateauing coverage and homogeneous mini-myotube templates. Coverage was normalized to the average of each user’s three most recent seeding attempts at expert level to account for cell line-to-cell line differences in absolute fiber coverage.

**Supplementary Figure 3.**
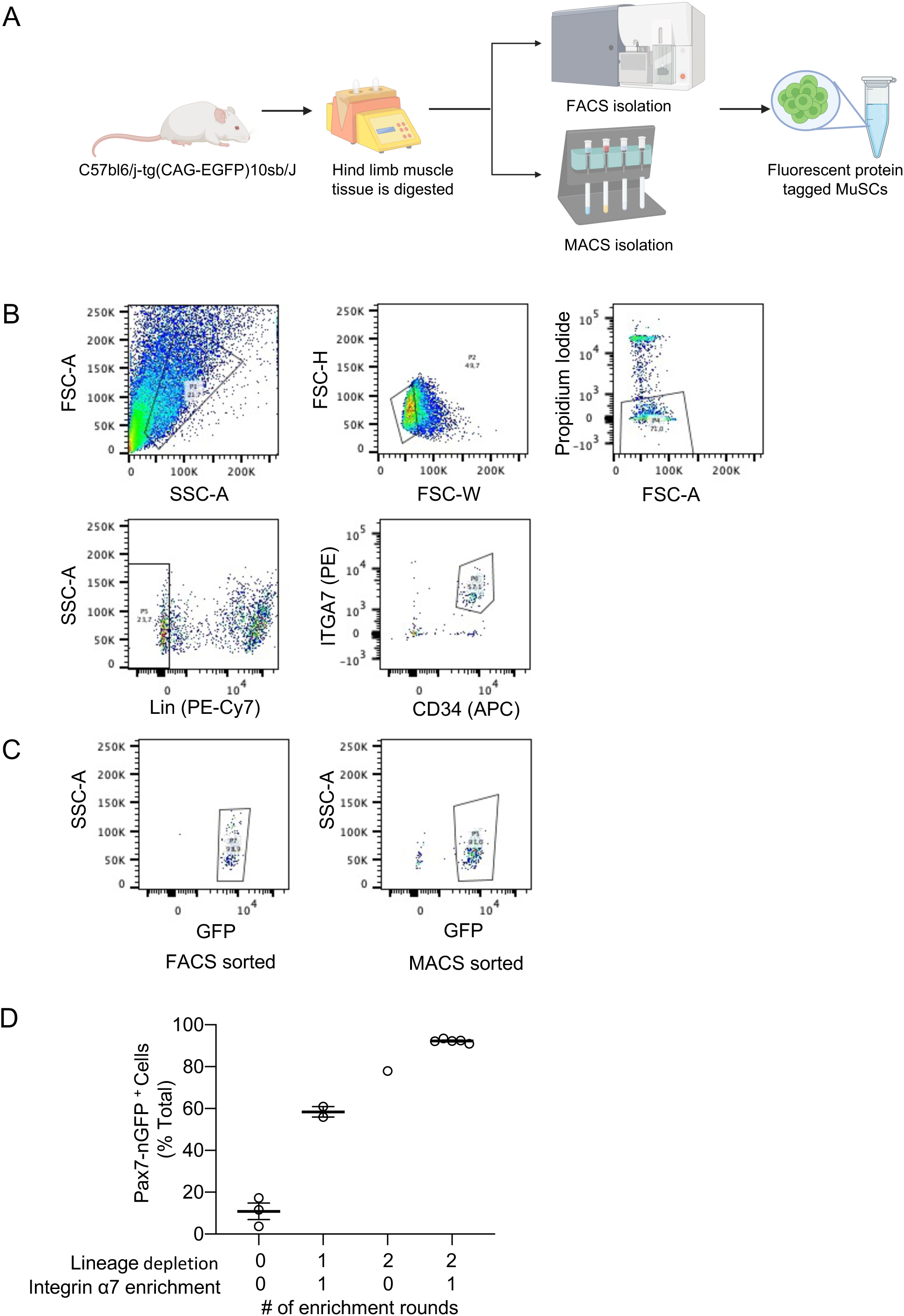
Gating strategy used for MACS or FACS based MuSCs isolation. **(A)** Schematic showcasing the workflow of MACS vs FACS MuSC isolation protocols. **(B)** Flow cytometry plots demonstrating gating strategy to enrich MuSCs from Pax7-nGFP transgenic mice using FACS. **(C)** Representative plots of green fluorescent protein (GFP)^+^ cells present in MuSC preparations derived from Pax7-nGFP tissue samples using FACS (left) as compared to MACS (right)**. (D)** Enumeration of % total GFP^+^ cells enriched from Pax7-nGFP tissue samples using a variety of optimisations that include the number of lineage depletion column rounds with or without an integin-α7 column enrichment round performed during the MACS protocol.

**Supplementary Figure 4.**
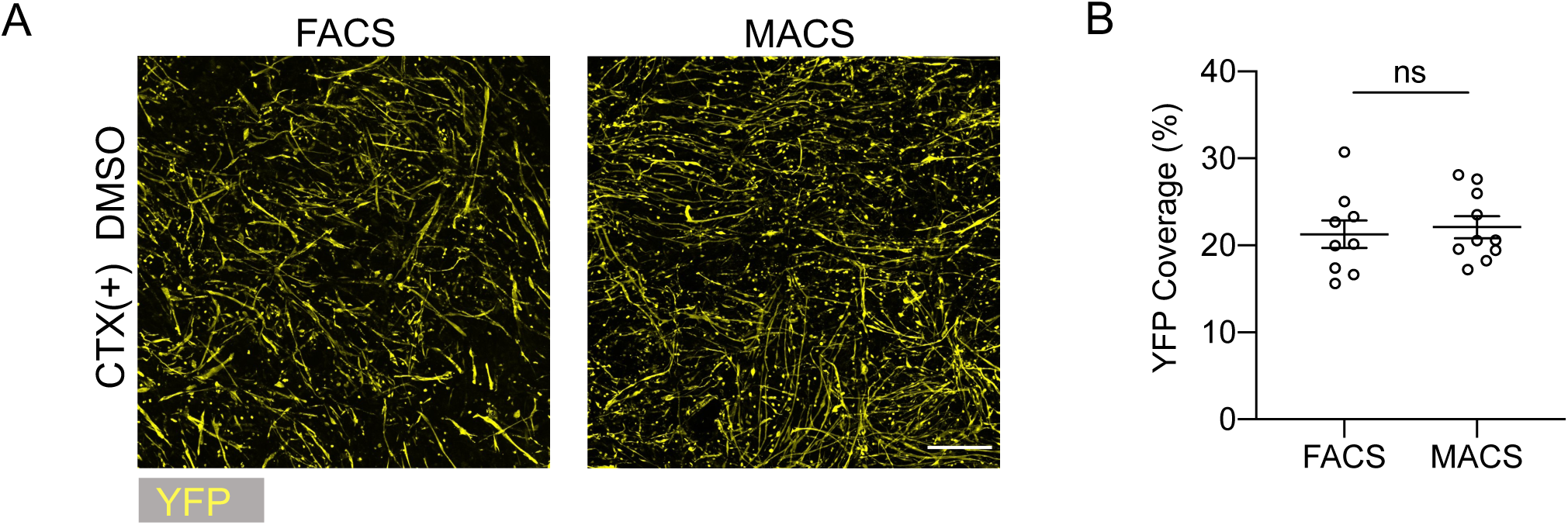
MACS and FACS sorted MuSCs are functionally similar in mini-MEndR assays. **(A)** Representative confocal images and **(B)** quantification of yellow fluorescent protein (YFP; yellow) immunostaining coverage of 18M mini-MEndR tissues at 10 DPI that were generated with FACS or MACS sorted MuSCs and treated with DMSO. Scale bar, 500 µm. Graphs display mean ± s.e.m.; unpaired two-tailed T-test with Welch’s correction, ns = no significance. n=9-10 tissues from N = 3 independent experiments per cell line.

## Notes

### Competing Interest Statement

The authors have declared no competing interest.

### Summary of Updates

-Title was edited to adjust focus. -A duplicated image was detected in Figure 3B, and upon thorough review of data, was determined to be a figure placeholder image that has now been updated with the phenotypic data that was generated for the condition (i.e. Day 14, 19F). -The manuscript text was edited to improve clarity.

## References

1. Brunet, A.; Goodell, M. A.; Rando, T. A. Ageing and Rejuvenation of Tissue Stem Cells and Their Niches. Nat. Rev. Mol. Cell Biol. 2023, 24 (1), 45–62. 10.1038/s41580-022-00510-w.

2. Dumont, N. A.; Wang, Y. X.; Von Maltzahn, J.; Pasut, A.; Bentzinger, C. F.; Brun, C. E.; Rudnicki, M. A. Dystrophin Expression in Muscle Stem Cells Regulates Their Polarity and Asymmetric Division. Nat. Med. 2015, 21 (12), 1455–1463. 10.1038/nm.3990.

3. Wang, Y. X.; Feige, P.; Brun, C. E.; Hekmatnejad, B.; Dumont, N. A.; Renaud, J. M.; Faulkes, S.; Guindon, D. E.; Rudnicki, M. A. EGFR-Aurka Signaling Rescues Polarity and Regeneration Defects in Dystrophin-Deficient Muscle Stem Cells by Increasing Asymmetric Divisions. Cell Stem Cell 2019, 24 (3), 419–432.e6. 10.1016/j.stem.2019.01.002.

4. Rozo, M.; Li, L.; Fan, C. M. Targeting Β1-Integrin Signaling Enhances Regeneration in Aged and Dystrophic Muscle in Mice. Nat. Med. 2016, 22 (8), 889–896. 10.1038/nm.4116.

5. Porpiglia, E.; Mai, T.; Kraft, P.; Holbrook, C. A.; de Morree, A.; Gonzalez, V. D.; Hilgendorf, K. I.; Frésard, L.; Trejo, A.; Bhimaraju, S.; Jackson, P. K.; Fantl, W. J.; Blau, H. M. Elevated CD47 Is a Hallmark of Dysfunctional Aged Muscle Stem Cells That Can Be Targeted to Augment Regeneration. Cell Stem Cell 2022, 29 (12), 1653–1668.e8. 10.1016/j.stem.2022.10.009.

6. Benjamin, D. I.; Both, P.; Benjamin, J. S.; Nutter, C. W.; Tan, J. H.; Kang, J.; Machado, L.; Klein, J. D. D.; de Morree, A.; Kim, S.; Liu, L.; Dulay, H.; Feraboli, L.; Louie, S. M.; Nomura, D. K.; Rando, T. A. Fasting Induces a Highly Resilient Deep Quiescent State in Muscle Stem Cells via Ketone Body Signaling. Cell Metab. 2022, 34 (6), 902–918.e6. 10.1016/j.cmet.2022.04.012.

7. Gilbert, P. M.; Havenstrite, K. L.; Magnusson, K. E. G.; Sacco, A.; Leonardi, N. A.; Kraft, P.; Nguyen, N. K.; Thrun, S.; Lutolf, M. P.; Blau, H. M. Substrate Elasticity Regulates Skeletal Muscle Stem Cell Self-Renewal in Culture. Science 2010, 329 (5995), 1078–1081. 10.1126/science.1191035.

8. Cosgrove, B. D.; Gilbert, P. M.; Porpiglia, E.; Mourkioti, F.; Lee, S. P.; Corbel, S. Y.; Llewellyn, M. E.; Delp, S. L.; Blau, H. M. Rejuvenation of the Muscle Stem Cell Population Restores Strength to Injured Aged Muscles. Nat. Med. 2014, 20 (3), 255–264. 10.1038/nm.3464.

9. Sacco, A.; Doyonnas, R.; Kraft, P.; Vitorovic, S.; Blau, H. M. Self-Renewal and Expansion of Single Transplanted Muscle Stem Cells. Nature 2008, 456 (7221), 502–506. 10.1038/nature07384.

10. Davoudi, S.; Xu, B.; Jacques, E.; Cadavid, J. L.; McFee, M.; Chin, C.-Y.; Meysami, A.; Ebrahimi, M.; Bakooshli, M. A.; Tung, K.; Ahn, H.; Ginsberg, H. J.; McGuigan, A. P.; Gilbert, P. M. MEndR: An in Vitro Functional Assay to Predict in Vivo Muscle Stem Cell-Mediated Repair. Adv. Funct. Mater. 2021.

11. Fleming, J. W.; Capel, A. J.; Rimington, R. P.; Wheeler, P.; Leonard, A. N.; Bishop, N. C.; Davies, O. G.; Lewis, M. P. Bioengineered Human Skeletal Muscle Capable of Functional Regeneration. BMC Biol. 2020, 18 (1), 145. 10.1186/s12915-020-00884-3.

12. Wang, J.; Broer, T.; Chavez, T.; Zhou, C. J.; Tran, S.; Xiang, Y.; Khodabukus, A.; Diao, Y.; Bursac, N. Myoblast Deactivation within Engineered Human Skeletal Muscle Creates a Transcriptionally Heterogeneous Population of Quiescent Satellite-like Cells. Biomaterials 2022, 284, 121508. 10.1016/j.biomaterials.2022.121508.

13. Rodenhizer, D.; Dean, T.; Xu, B.; Cojocari, D.; McGuigan, A. P. A Three-Dimensional Engineered Heterogeneous Tumor Model for Assessing Cellular Environment and Response. Nat. Protoc. 2018, 13 (9), 1917–1957. 10.1038/s41596-018-0022-9.

14. Sambasivan, R.; Gayraud-Morel, B.; Dumas, G.; Cimper, C.; Paisant, S.; Kelly, R. G.; Tajbakhsh, S. Distinct Regulatory Cascades Govern Extraocular and Pharyngeal Arch Muscle Progenitor Cell Fates. Dev Cell 2009, 16 (6), 810–821. 10.1016/j.devcel.2009.05.008S1534-5807(09)00210-X [pii].

15. Davoudi, S.; Chin, C.-Y.; Cooke, M. J.; Tam, R. Y.; Shoichet, M. S.; Gilbert, P. M. Muscle Stem Cell Intramuscular Delivery within Hyaluronan Methylcellulose Improves Engraftment Efficiency and Dispersion. Biomaterials 2018, 173, 34–46. 10.1016/j.biomaterials.2018.04.048.

16. Laumonier, T.; Bermont, F.; Hoffmeyer, P.; Kindler, V.; Menetrey, J. Human Myogenic Reserve Cells Are Quiescent Stem Cells That Contribute to Muscle Regeneration after Intramuscular Transplantation in Immunodeficient Mice. Sci. Rep. 2017, 7 (1), 3462. 10.1038/s41598-017-03703-y.

17. Hardy, D.; Besnard, A.; Latil, M.; Jouvion, G.; Briand, D.; Thépenier, C.; Pascal, Q.; Guguin, A.; Gayraud-Morel, B.; Cavaillon, J. M.; Tajbakhsh, S.; Rocheteau, P.; Chrétien, F. Comparative Study of Injury Models for Studying Muscle Regeneration in Mice. PloS One 2016, 11 (1), e0147198. 10.1371/journal.pone.0147198.

18. Giordani, L.; He, G. J.; Negroni, E.; Sakai, H.; Law, J. Y. C.; Siu, M. M.; Wan, R.; Corneau, A.; Tajbakhsh, S.; Cheung, T. H.; Le Grand, F. High-Dimensional Single-Cell Cartography Reveals Novel Skeletal Muscle-Resident Cell Populations. Mol. Cell 2019, 74 (3), 609–621.e6. 10.1016/J.MOLCEL.2019.02.026.

19. Jacques, E.; Kuang, Y.; Kann, A. P.; Le Grand, F.; Krauss, R. S.; Gilbert, P. M. The Mini-IDLE 3D Biomimetic Culture Assay Enables Interrogation of Mechanisms Governing Muscle Stem Cell Quiescence and Niche Repopulation. eLife 2022, 11, e81738. 10.7554/eLife.81738.

20. Li, N. T.; Wu, N. C.; Cao, R.; Cadavid, J. L.; Latour, S.; Lu, X.; Zhu, Y.; Mijalkovic, M.; Roozitalab, R.; Landon-Brace, N.; Notta, F.; McGuigan, A. P. An Off-the-Shelf Multi-Well Scaffold-Supported Platform for Tumour Organoid-Based Tissues. Biomaterials 2022, 291, 121883. 10.1016/j.biomaterials.2022.121883.

21. Cao, R.; Li, N. T.; Cadavid, J. L.; Latour, S.; Tan, C. M.; McGuigan, A. P. An Automation Workflow for High-Throughput Manufacturing and Analysis of Scaffold-Supported 3D Tissue Arrays; preprint; Bioengineering, 2022. 10.1101/2022.08.20.504600.

